# Heartbeat-Related Bodily Processing Shapes Transition Patterns in Self-Related Spontaneous Thought

**DOI:** 10.64898/2026.08.25.747022

**Authors:** Mai Sakuragi, Kazushi Shinagawa, Yuri Terasawa, Yuto Tanaka, Satoshi Umeda

**Affiliations:** Department of Psychology, Keio University, 2-15-45 Mita, Minato-ku, Tokyo, 108-8345, Japan; Japan Society for the Promotion of Science, Kojimachi Business Center Building, 5-3-1, Kojimachi, Chiyoda-ku, Tokyo 102-0083, Japan; Department of Information Medicine, National Institute of Neuroscience, National Center of Neurology and Psychiatry, Tokyo 187-8502, Japan; Keio University Global Research Institute, 2-15-45 Mita, Minato-ku, Tokyo, 108-8345, Japan

**Keywords:** Spontaneous Thought, Autonomic Nervous Activity, Interoception

## Abstract

Spontaneous thought changes over time, yet the moment-to-moment factors shaping these changes remain poorly understood. We examined whether heartbeat-related bodily processing, operating largely outside explicit awareness, is associated with the organization of ongoing thought. Forty adults performed an auditory attention task with intermittent thought probes in which auditory events were scheduled either 200 ms after each detected R peak (synch condition) or independently of ongoing cardiac timing (asynch condition), with occasional omissions in both conditions. Heartbeat-synchronous omissions in this paradigm have been shown to induce cardiac deceleration and modulate heartbeat-evoked potentials (HEPs). The score of the heartbeat counting task (HCT) served as a behavioral index related to cardiac interoceptive accuracy. The results showed that higher HCT score was associated with a stronger synch–asynch shift toward more self-related and less task/self-unrelated thought. HEPs showed a synch-related negative shift across several thought groups, although the magnitude of this condition effect did not reliably differ among thought groups. Overall thought-group distributions were similar across conditions, while transition analyses suggested a tendency in the synch condition toward more frequent transitions linking predominantly interoceptive stimulus-dependent thought with both on-task and self-related thought. Together, these findings suggest that heartbeat-related bodily processing may influence the organization of spontaneous thought, particularly in individuals with greater cardiac interoceptive accuracy. Bodily signals may therefore act as an automatic constraint on the ongoing stream of thought, biasing which thought contents and transitions become more likely over time.

## Introduction

Spontaneous thought has been conceptualized as mental activity whose generation and progression are relatively weakly constrained by deliberate control, such as explicit goals and immediate task demands (Christoff et al., 2016; Smallwood & Schooler, 2015). Research in this field has traditionally focused on quantifying the frequency of task-unrelated thoughts and classifying their content, including dimensions such as task-relatedness, stimulus-dependency, temporal orientation, intentionality, and self-relatedness (Babo-Rebelo et al., 2016; Kane et al., 2017; Seli et al., 2016; Smallwood et al., 2009, 2011; Smallwood & Schooler, 2006; Stawarczyk et al., 2011). More recently, attention has shifted toward the temporal dynamics of ongoing thought—how mental states persist and transition over time (Christoff et al., 2016; Mittner et al., 2014). Methodological approaches that estimate transitions and temporal occupancy among thought states have made it possible to characterize the organization of thought as a dynamic process rather than as a collection of isolated reports (Shinagawa et al., 2023; Zanesco et al., 2020).

Although temporal approaches have clarified how thought states persist and transition over time, the factors underlying these dynamics remain incompletely understood. Existing accounts have emphasized current concerns and goal-related thought generation, executive control over task focus, and meta-awareness of attentional drift (Klinger, 2013; McVay & Kane, 2010; Schooler et al., 2011). However, many spontaneous shifts in thought occur without an identifiable trigger; indeed, approximately half of everyday mind-wandering episodes have been reported without a clear precipitating event (Faber & D’Mello, 2018). This leaves open whether continuously varying state factors, operating on a similar timescale to thought transitions and often outside conscious awareness or voluntary control, contribute to whether ongoing thought persists or shifts.

Among such state factors, autonomic cardiac activity is a particularly plausible candidate because it varies continuously, often outside conscious awareness, and is tightly coupled with central neural processing. In line with the neurovisceral integration framework, central and peripheral processes are thought to interact bidirectionally, and indices such as heart rate variability (HRV) have been linked to regulatory flexibility (Thayer et al., 2009, 2012). Conversely, lower vagally mediated HRV, including lower RMSSD, has been associated with more rigid and perseverative cognitive tendencies such as worry and rumination (Ottaviani et al., 2015, 2017). Although these findings do not directly address spontaneous-thought transitions, they suggest that autonomic state is related to the flexibility versus persistence of ongoing cognition. This raises the possibility that moment-to-moment autonomic state may contribute to the automatic constraints that bias whether ongoing thought persists or shifts over time.

A plausible route through which autonomic state could influence ongoing thought is interoceptive processing, whereby the nervous system senses and represents signals arising from within the body (Khalsa et al., 2018). This processing involves regions including the insula and is closely linked to central autonomic regulation (Craig, 2002; Critchley & Harrison, 2013; Menon & Uddin, 2010). In other words, changes in bodily state are continuously registered by the brain and may therefore influence which kinds of thoughts become more likely as cognition unfolds. This possibility is particularly relevant to self-related cognition. Cardiac and other interoceptive signals contribute to the representation of one’s own bodily state and have been linked to bodily self-consciousness. Heartbeat-evoked potentials (HEPs), electrophysiological responses time-locked to individual heartbeats and commonly used as an index of cortical processing of cardiac signals, have also been shown to vary with the self-relatedness of spontaneous thought (Babo-Rebelo et al., 2016; Park et al., 2014; Tallon-Baudry et al., 2018). These findings suggest that changes in heartbeat-related bodily processing may bias ongoing thought toward or away from self-related content, providing one possible route by which bodily state contributes to the organization of spontaneous thought.

This possibility is consistent with our previous work linking autonomic activity and individual differences in cardiac interoception to self-related thought. One behavioral dimension of cardiac interoception is cardiac interoceptive accuracy, which refers to the accuracy of performance on tasks designed to assess the perception of one’s own cardiac activity (Garfinkel et al., 2015). In our previous and present studies, this was assessed using the heartbeat counting task (HCT; Schandry, 1981), with HCT performance used as a behavioral index related to cardiac interoceptive accuracy. We found that lower HRV was more strongly associated with the persistence of self-related thought over time among individuals with higher heartbeat-counting performance (Sakuragi, Shinagawa, et al., 2024; Sakuragi & Umeda, 2025). We also found that experimentally induced heart-rate changes were associated with more frequent self-related thought among individuals with higher heartbeat-counting performance (Sakuragi et al., 2023). However, these studies did not establish how experimentally modulating cardiac state was accompanied by changes in the cortical processing of cardiac signals or in the temporal organization of ongoing thought. The experimentally induced cardiac changes also varied substantially across individuals, limiting control over the physiological manipulation (Sakuragi et al., 2023; Sakuragi, Tanaka, et al., 2024). A further step is therefore to manipulate heartbeat-related bodily processing under more controlled conditions while simultaneously characterizing spontaneous-thought organization and heartbeat-related cortical activity.

The heartbeat-synchronous auditory omission paradigm provides a candidate method for experimentally probing the link between cardiac activity and ongoing thought. In this paradigm, sounds are presented in temporal relation to the heartbeat, and occasional omissions perturb the coupling between cardiac timing and external auditory events. Previous cardio-auditory omission studies have shown that such omissions modulate HEP and cardiac responses, including heart-rate deceleration (Banellis & Cruse, 2020; Pelentritou et al., 2024, 2025). In a validation study using an active auditory detection task closely matching the present setting, we further showed that heartbeat-synchronous omissions were accompanied by RR-interval prolongation and enhanced HEPs, with only minor effects on behavioral performance (Sakuragi et al., 2026). Thus, this paradigm provides a task-embedded means of modulating cardiac responses and heartbeat-related cortical processing without requiring participants to explicitly attend to or become aware of the heartbeat–sound relationship.

The present study applied this paradigm to examine whether covert heartbeat-related bodily processing was associated with changes in the distribution and transition patterns of spontaneous thought. Participants performed an auditory detection task in which sounds were presented either at a fixed latency from the heartbeat (synch condition) or independently of ongoing cardiac timing (asynch condition). In both conditions, sounds were occasionally omitted. In the synch condition, these omissions occurred within a heartbeat-locked auditory context and have been shown in this task to be accompanied by cardiac deceleration and enhanced HEPs (Sakuragi et al., 2026).

Accordingly, we tested three hypotheses concerning thought organization, heartbeat-related cortical processing, and individual differences in cardiac interoception. First, if heartbeat-related bodily processing contributes to ongoing thought dynamics, we expected the distribution and transition patterns of thought states to differ between the synch and asynch conditions, with differences particularly involving self-related thought. Second, given previous evidence that HEPs vary with the self-relatedness of spontaneous thought, we expected self-related thought to be accompanied by enhanced HEP responses. Third, based on previous findings that associations between cardiac state and self-related thought are stronger in individuals with higher HCT performance, we expected condition-related differences in thought distribution to vary with HCT performance, such that participants with higher HCT performance would show a greater relative occurrence of self-related thought in the synch condition.

## Methods and Materials

### Participants

Forty-two healthy undergraduate students, graduate students, and working adults participated in the experiment (19 males, 23 females; mean age = 22.2 years; age range = 18–30 years). All participants reported normal hearing and had normal or corrected-to-normal vision. The present study analyzed the same participant sample as the companion physiological study of the heartbeat-synchronous auditory omission paradigm (Sakuragi et al., 2026). The sample size was determined a priori for the physiological manipulation analysis reported in that study, using Bayes factor design analysis based on omission-evoked cardiac responses; full details of the sample-size planning are provided in Sakuragi et al. (2026). The resulting target sample size was N = 42. Two participants were excluded from the analysis due to excessive noise in the ECG recordings; therefore, the final analytic sample comprised 40 participants (18 males, 22 females; mean age = 22.3 years; age range = 18–30 years).

Before participation, we screened participants for factors that could affect physiological recordings (e.g., EEG and ECG) or performance on the psychological task. The exclusion criteria were as follows: history of epileptic seizures; history of surgical procedures involving the head; cardiac disease; current use of medications affecting psychiatric or neurological functions, analgesics, cold medicine, or anti-allergic agents; difficulty understanding the experimental instructions in Japanese; and consumption of vasoactive substances, including food or drinks containing caffeine, alcohol, or nicotine, within three hours before the experiment.

Participants wearing makeup, particularly around the eyes, were asked to report this to the experimenter on the day of the experiment and to remove it if necessary. If they reported being unable to comply with this requirement, they were not allowed to participate. Whether participants met any of these criteria was confirmed verbally on the day of the experiment. No participants were excluded based on these criteria after enrollment. This study was approved by the Keio University Research Ethics Committee (Approval No. 240040000) and was conducted in accordance with the Declaration of Helsinki.

### Apparatus

Throughout the experimental task, ECG was recorded using an MP-150 system and AcqKnowledge software (Biopac Systems, Santa Barbara, CA). ECG was recorded in a three-lead configuration, with electrodes placed on the right hand and both feet. ECG signals were used for online R-wave detection, scheduling heartbeat-synch auditory events, calculating RR intervals, and time-locking heartbeat-evoked potentials. EEG was recorded simultaneously with ECG using NetStation 5.3.0.1 and a 64-channel HydroCel Geodesic Sensor Net. In the present paper, we report ECG- and EEG-based measures relevant to heartbeat-synch stimulation, thought-state dynamics, and heartbeat-evoked potentials.

### Procedure

#### Heartbeat counting task (HCT)

Cardiac interoceptive accuracy was assessed using the Heartbeat Counting Task (HCT; Schandry, 1981). In the present study, HCT performance was used as a behavioral index related to cardiac interoceptive accuracy. The task included a resting ECG measurement, heartbeat-counting trials, and a time-estimation task. The time estimation task was administered to discourage participants from relying on perceived elapsed time when estimating heartbeat counts; however, only HCT performance was analyzed in the present paper. During the task, participants used their left hand to enter the counted heartbeats and confidence ratings on the keyboard, while their right hand rested on their lap. In each heartbeat-counting trial, participants silently counted their heartbeats for a predefined interval and reported the number of heartbeats. They were instructed not to count heartbeats by touching their body, pressing against the backrest or desk, or holding their breath. Following the revised HCT instructions (Desmedt et al., 2018), participants were also instructed not to guess heartbeats they could not feel and not to calculate the count based on prior knowledge of their heart rate or perceived elapsed time. After each trial, participants rated their confidence in their performance on a scale from 1 to 10.

#### Heartbeat-synchronous auditory task

The main task was an auditory attention task with intermittent experience-sampling probes (Figure 1). Participants were instructed to press the “9” key with their left hand as quickly and accurately as possible whenever they heard a tone. Auditory stimuli were 1000-Hz pure tones presented at 65 dB for 100 ms.

**Figure 1.**
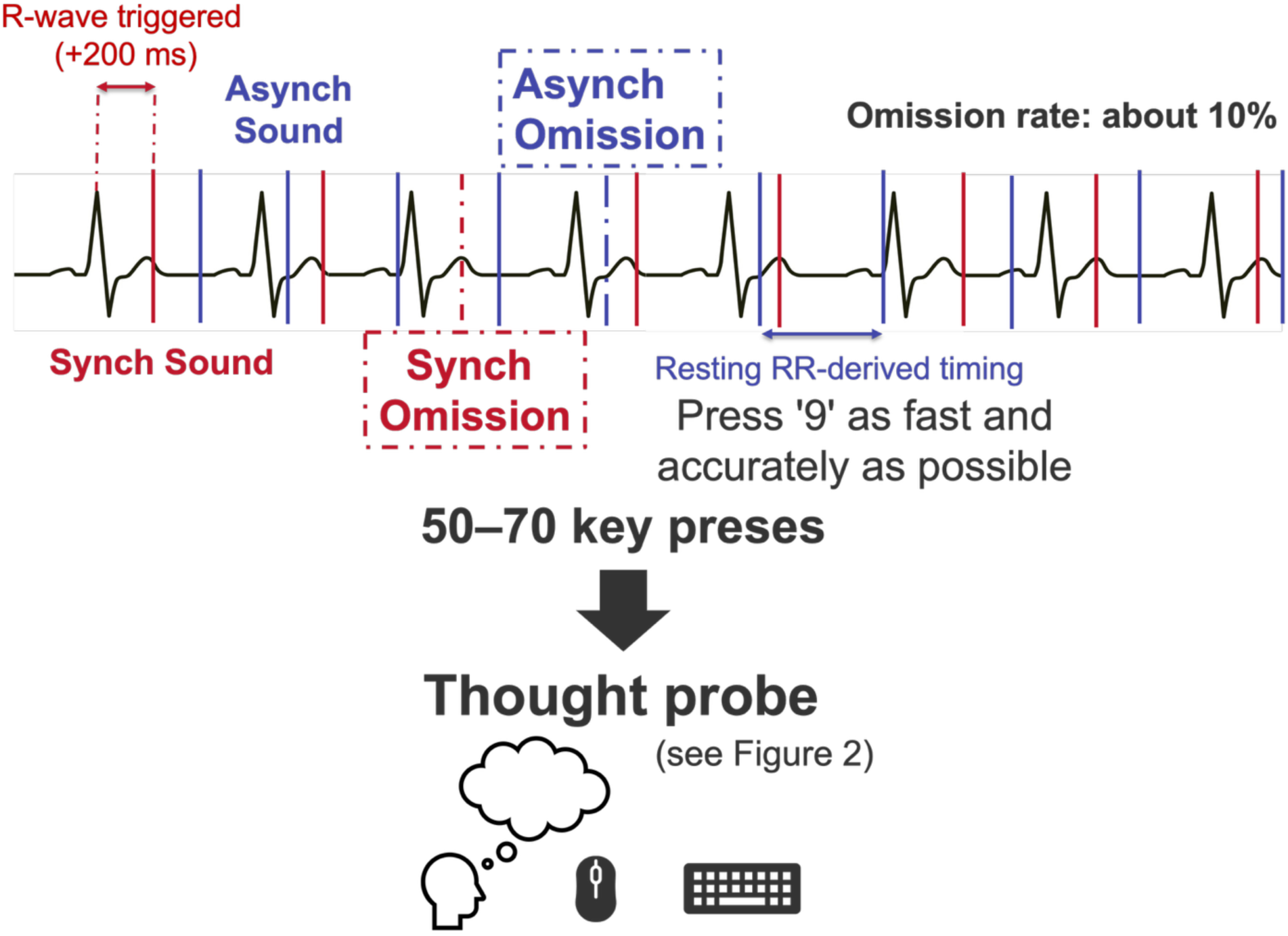
Experimental task procedure. Participants performed an auditory detection task in which they were instructed to press the “9” key as quickly and accurately as possible whenever a tone was presented. In the synch condition, auditory events were scheduled 200 ms after online detection of each R peak. In the asynch condition, events were scheduled according to a pseudorandom sequence with inter-event intervals drawn from each participant’s resting RR-interval range and were therefore independent of ongoing R waves. In both conditions, approximately 10% of auditory events were omissions. After 50–70 key presses, a thought probe was presented to assess the participant’s thought content during the immediately preceding period. Each segment ending with a thought probe constituted one trial, and participants completed 15 trials per condition. The detailed thought-probe procedure is shown in Figure 2. Red and blue markers schematically illustrate auditory-event timing in the separate synch and asynch conditions, respectively. Alt text. Schematic of the auditory attention task showing synch and asynch auditory-event timing relative to the ECG. In the synch condition, sounds and occasional omissions were scheduled 200 ms after detected R waves, whereas in the asynch condition they followed a pseudorandom timing sequence derived from each participant’s resting RR intervals and were independent of ongoing R waves; omissions occurred on approximately 10% of scheduled events in both conditions. Participants pressed the “9” key for each presented sound, and a thought probe was administered after 50–70 key presses.

Auditory events were presented in two blocks: a heartbeat-synchronous (synch) condition and a heartbeat-asynchronous (asynch) condition. Before the task, five minutes of resting physiological data were recorded without auditory stimulation to estimate each participant’s resting RR-interval range and to generate the pseudorandom sound sequence for the asynch condition. In the synch condition, tones or omissions were scheduled 200 ms after online detection of the participant’s R wave. In the asynch condition, tones or omissions were scheduled according to a pre-generated pseudorandom sequence with inter-event intervals drawn from a uniform distribution between the minimum and maximum resting RR intervals of each participant. Thus, the asynch sequence approximated the participant’s resting cardiac rhythm while remaining independent of ongoing R waves. The order of the synch and asynch blocks was randomized across participants. In both conditions, 10% of auditory events were omissions, and at least three tone-present events were inserted between successive omissions.

The physiological effects of the heartbeat-synchronous auditory omission paradigm used here have been reported separately (Sakuragi et al., 2026). In that analysis, heartbeat-synchronous omissions induced cardiac deceleration and enhanced heartbeat-evoked potentials. In the present study, we used the same paradigm to test whether a heartbeat-synchronous auditory context, designed to modulate cardiac responses and heartbeat-related cortical processing, alters spontaneous-thought dynamics.

Thought probes were presented after every 50 to 70 key presses, with the exact interval randomly determined from a uniform distribution. Each segment ending with a thought probe was defined as one trial. Participants completed 15 trials in each condition, for a total of 30 trials. Thus, the number of scheduled thought probes was balanced across the synch and asynch conditions. At each probe, participants selected the thought category that best represented their mental content during the period immediately preceding the probe. They were instructed to select the content that had occupied the greatest portion of that period or, when multiple thoughts occurred, the content that left the strongest impression They selected one of eight categories: task-focus, task-related thought, exteroception, interoception, self-related thought, task/self-unrelated thought, mind-blanking, or forgetfulness. Participants responded by pressing the corresponding number key. Before the experiment, the experimenter provided oral definitions of all categories and instructed participants to report their ongoing experience as accurately and honestly as possible. Participants were told that physiological recordings, including EEG, could provide information relevant to their mental state and that clearly inconsistent or unreliable reports could result in data exclusion. They were also informed that task-unrelated thoughts, mind-blanking, and other response categories were all acceptable and that no particular category was considered preferable. As an additional quality-control criterion, participants were screened for an excessive concentration of Mind-blanking or Forgetfulness responses, which could indicate insufficient engagement with the task or unreliable thought reporting. Participants were to be excluded if such responses exceeded eight of the 15 thought probes in either experimental condition. No participant met this criterion.

When participants selected a category other than mind-blanking or forgetfulness, they subsequently rated the phenomenological features of the thought episode. These ratings included task concentration, temporal orientation, arousal, valence, self-relatedness, perspective, bodily information, contemplation, and intentionality. Continuous dimensions were assessed using visual analog scales, whereas temporal orientation, valence, perspective, and intentionality were assessed using categorical response options; the full set of response scales is summarized in Table 1. Some probes were skipped when the initial category response determined their values. Specifically, when participants selected task-focus, temporal orientation was coded as “now”; when they selected task/self-unrelated thought, self-relatedness and bodily information were coded as 0. When participants selected mind-blanking or forgetfulness, all subsequent phenomenological probes were skipped and coded as a no-response category. The probe sequence and category-dependent skipping rules are illustrated in Figure 2.

**Figure 2.**
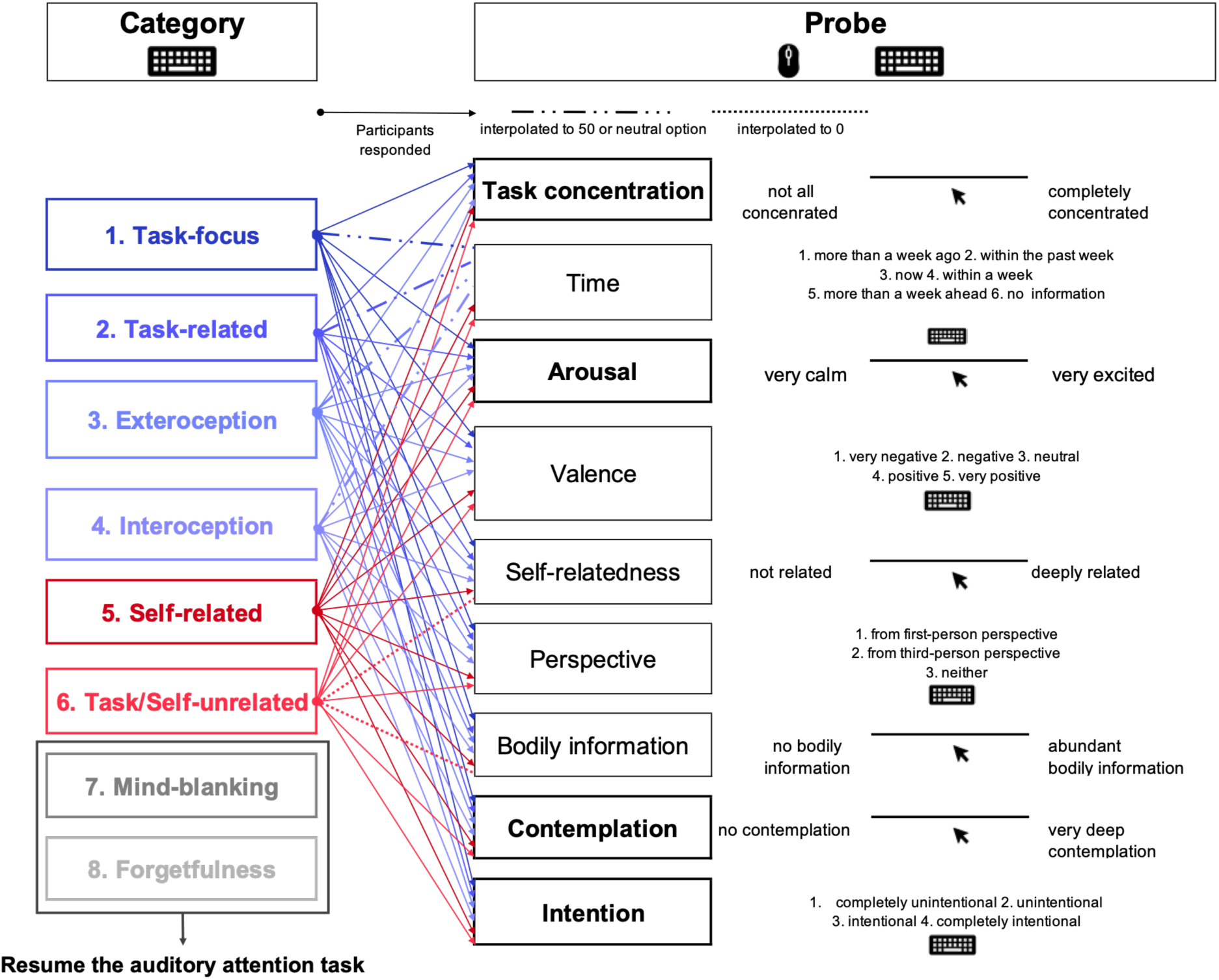
Thought categories selected by participants and their corresponding probes. Colors indicate the broad relationship of each response category to the ongoing task and sensory events: blue denotes thoughts related either to the task itself or to sensory information arising during the task, red denotes task-unrelated and stimulus-independent thoughts, and gray denotes the absence of reportable thought content. Participants first selected one of eight categories that best represented their thought content during the period immediately preceding the probe, selecting the content that occupied the greatest portion of that period or left the strongest impression. Subsequently, based on the selected category, questions regarding various components of the thought (connected to each category by arrows) were presented. Participants responded to the probes either by clicking the appropriate position on a solid line with their left hand using a mouse or by entering the number of the most applicable option from the choices presented on the screen. Scores on the solid line were quantified with 0 at the left end, 50 at the center, and 100 at the right end. If “Mind-blanking” or “Forgetting” was selected, the thought probe session terminated, and the auditory attention task resumed. Probes indicated in bold and enclosed in thick borders were consistently presented whenever any category other than “Mind-blanking” or “Forgetting” was selected. Questions regarding components that were self-evident from each category’s definition were not presented during the task; instead, appropriate values were interpolated during subsequent data processing. Probes connected to category labels by dashed lines were interpolated with a value of 50, while those connected by dotted lines were interpolated with a value of 0. If “Mind-blanking” or “Forgetting” was selected, responses to questions regarding thought components were coded as a separate “no response” category for use in subsequent analyses. Alt text. Flow diagram of the thought-probe procedure. Participants first selected one of eight thought categories: Task-focus, Task-related, Exteroception, Interoception, Self-related, Task/Self-unrelated, Mind-blanking, or Forgetfulness. The first six categories led to category-dependent follow-up questions assessing phenomenological features such as task concentration, temporal orientation, arousal, valence, self-relatedness, perspective, bodily information, contemplation, and intentionality. Some follow-up values were predetermined from the selected category and were therefore not directly queried. Selecting Mind-blanking or Forgetfulness terminated the probe sequence without further phenomenological questions.

**Table 1.** Phenomenological probe items and response scales.

| Probe item | Scale / options |
| --- | --- |
| Task concentration | 0 = not at all concentrated; 100 = completely concentrated |
| Temporal orientation | 1 = more than a week ago; 2 = within the past week;<br>3 = now; 4 = within a week; 5 = more than a week ahead; 6 = no temporal information |
| Arousal | 0 = very calm; 100 = very excited |
| Valence | 1 = very negative; 2 = negative; 3 = neutral;<br>4 = positive; 5 = very positive |
| Self-relatedness | 0 = not related to me; 100 = deeply related to me |
| Perspective | 1 = first-person perspective; 2 = third-person perspective;<br>3 = neither |
| Bodily information | 0 = no bodily information; 100 = abundant bodily information |
| Contemplation | 0 = no contemplation; 100 = very deep contemplation |
| Intentionality | 1 = completely unintentional; 2 = unintentional;<br>3 = intentional; 4 = completely intentional |

The duration of each trial varied depending on the participant’s resting RR interval and response speed to the thought probes, but typically ranged from 1.5 to 2 min. The entire task lasted approximately 25–30 min. After the task, participants completed the STAI-State questionnaire again and an original questionnaire assessing their daily thought patterns during the preceding week.

### Data Preprocessing

#### ECG data

R-peaks were detected from the raw ECG waveforms using AcqKnowledge. To exclude artifactual RR intervals, threshold-based artifact rejection was applied separately to the RR intervals obtained during the heartbeat counting task and the auditory attention task for each participant, using the median RR interval within each task as the reference. RR intervals exceeding 1.5 times the task-specific median RR interval or falling below two-thirds of the task-specific median RR interval were defined as outliers and excluded from subsequent processing. Physiological data were synchronized with the heartbeat counting task and the auditory attention task using event triggers recorded in the experimental control logs. To control for the influence of motor responses and transient cognitive load associated with thought-probe responses, intervals from probe presentation to response completion were identified and excluded, and only RR data recorded during task performance were extracted.

#### EEG data

EEG data preprocessing was performed using the EEGLAB toolbox (Delorme & Makeig, 2004) and its MATLAB plugins (MathWorks, USA). Raw EEG data were downsampled to 250 Hz, and a band-pass filter was applied from 0.5 Hz to 40 Hz. Artifact Subspace Reconstruction (ASR) was used to remove and correct artifacts. First, bad channels were detected (correlation with adjacent channels < 0.85; high-frequency noise *SD* > 4) and interpolated using spherical splines. Subsequently, ASR correction was applied to intervals containing short-duration burst noise (Burst Criterion = 30 *SD*). After ASR, the data were re-referenced to the common average. Independent component analysis using the extended Infomax algorithm was performed to remove biological artifacts (eye movements, muscle activity, ECG). ICLabel (Pion-Tonachini et al., 2019) was used to classify independent components automatically. Components exceeding the following probability thresholds were excluded: Eye movements > 0.85, Muscle activity > 0.85, ECG > 0.60, Line noise > 0.85, and Channel noise > 0.85. Finally, a second strict cleaning (Burst Criterion = 15 *SD*, Window Criterion = 0.2) was performed to remove remaining significant artifacts.

### Statistical Analysis

#### HCT performance

Performance scores for the HCT were calculated for each trial using the formula:

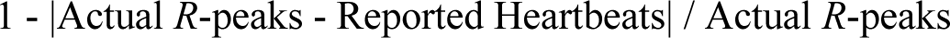

The mean score across six trials was used as the HCT performance score and served as a behavioral index related to cardiac interoceptive accuracy.

#### Classification of Thought Groups

Based on the framework proposed by Stawarczyk et al. (2011), we recategorized the thought probe responses into five groups along two primary axes: task-relatedness and stimulus-independence (Figure 3). Given that the central focus of this study is the transition from external task engagement to internal thought, the categories were grouped as follows:

- Group 1 (On-task): Combined Task-focus and Task-related categories, representing states oriented toward the experimental task.
- Group 2 (Stimulus-dependent): Combined Exteroception and Interoception. These are thoughts unrelated to the task itself that are triggered by sensory input from either the external environment or the body.
- Group 3 (Self-related): Represented by Self-related thoughts. These are task-unrelated and stimulus-independent, which we identified as a critical axis, given prior literature emphasizing the role of self-relevance in spontaneous cognition.
- Group 4 (Task/Self-unrelated): Represented by Task/Self-unrelated thoughts (e.g., general knowledge), which are stimulus-independent but lack personal relevance.
- Group 5 (Thought absence): Combined Mind-blanking and Forgetfulness, representing a lack of reportable thought content.

**Figure 3.**
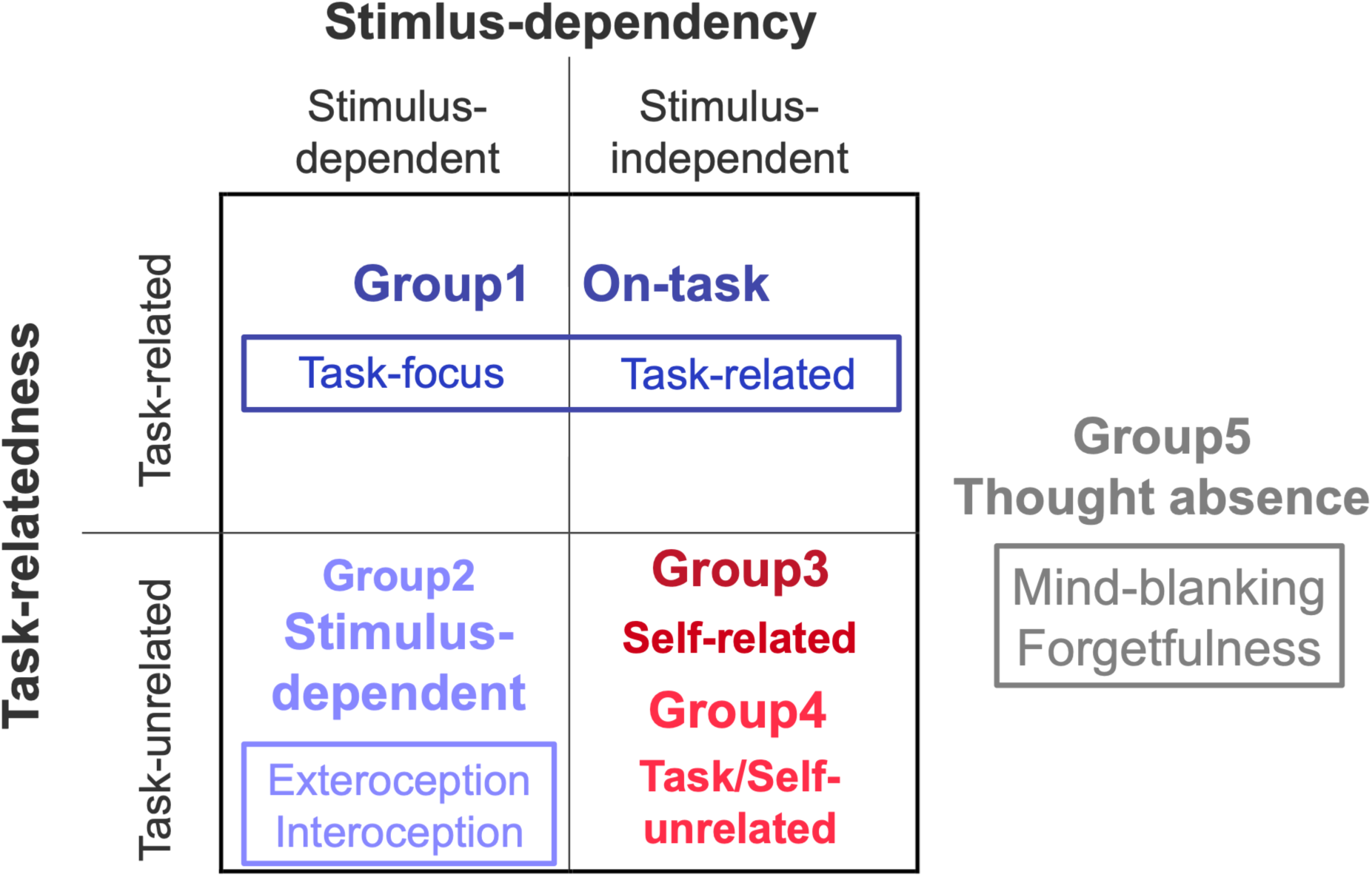
Classification of thought groups based on task-relatedness and stimulus-independence. The vertical axis represents task-relatedness (task-related vs. task-unrelated), and the horizontal axis represents stimulus-independence (stimulus-dependent vs. stimulus-independent). The four central quadrants illustrate the classification of thought categories according to the model of Stawarczyk et al. (2011). The colored boxes and labels correspond to the five thought groups defined in this study: • Group 1 (Blue): Encompasses Task-focus and Task-related categories. • Group 2 (Light Blue): Includes Exteroception and Interoception, representing thoughts triggered by sensory inputs from the environment or the body. • Group 3 (Dark Red): Represents Self-related thoughts, a subset of stimulus-independent, task-unrelated thoughts categorized by self-relatedness. • Group 4 (Red): Represents Task/Self-unrelated thoughts. • Group 5 (Gray): Shown to the right of the main quadrants, represents Mind-blanking and Forgetfulness, indicating an absence of reportable thought content. Alt text: Conceptual classification of five thought groups according to task-relatedness and stimulus-dependency. Group 1, On-task thought, occupies the task-related region and includes Task-focus and Task-related thoughts. Group 2, Stimulus-dependent thought, occupies the task-unrelated and stimulus-dependent region and includes Exteroception and Interoception. Groups 3 and 4 occupy the task-unrelated and stimulus-independent region: Group 3 represents Self-related thought, whereas Group 4 represents Task/Self-unrelated thought. Group 5, Thought absence, is shown outside the two-dimensional classification and includes Mind-blanking and Forgetfulness.

#### Statistical Analysis of Thought Groups Subjective Characteristics of Thought Groups

To characterize the subjective profiles of the thought groups, we performed Bayesian analyses of the continuous and categorical phenomenological probe ratings. Because the phenomenological probes were not administered following Mind-blanking or Forgetfulness responses, Group 5 (Thought absence) was excluded from these analyses. Analyses of phenomenological ratings therefore included Groups 1–4 and were conducted separately for the synch and asynch conditions.

- Continuous Probes: For probes rated on a continuous scale (Task concentration, Arousal, Self-relatedness, Bodily information, and Contemplation; Probes 1, 3, 5, 7, and 8), thought group was included as the sole fixed effect to examine differences in subjective ratings across Groups 1–4.
- Categorical Probes: For categorical probes (Time, Valence, Perspective, and Intention; Probes 2, 4, 6, and 9), the report probability of each response option was calculated from the frequency with which that option was selected within each participant and thought group. The interaction between thought group and response option was included as the fixed effect to examine whether the distribution of responses differed across Groups 1–4.

#### Objective Behavioral and Cardiac Characteristics of Thought Groups

To characterize the thought groups using objective behavioral and cardiac measures, we examined three measures of behavioral variability, response performance, and cardiac activity. Unlike the phenomenological ratings, these measures were available for Group 5 (Thought absence); therefore, all five thought groups were included in these analyses.

- Reaction-Time Coefficient of Variation: RTCV was calculated as the standard deviation of reaction times divided by the mean reaction time, providing an index of trial-to-trial variability in behavioral responses.
- Response Error Rate: A response error was defined as a tone event for which no response occurred before the subsequent tone event. For each trial, the error rate was calculated as the number of such errors divided by the total number of tone events.
- Standardized RR Interval: RR intervals were derived from consecutive R peaks in the ECG and standardized within participant using values across the entire experiment. Epochs immediately following sound omissions were excluded because omissions were known to evoke transient changes in RR interval (Sakuragi et al., 2026).

For each measure, condition, thought group, and their interaction were included as fixed effects to examine whether the objective measures differed across thought groups and whether these differences varied between the synch and asynch conditions. Based on the fitted models, follow-up posterior contrasts were calculated where relevant. For response errors, posterior-predicted error probabilities were estimated for each condition and thought group, and synch–asynch contrasts were calculated separately within each thought group. For standardized RR interval, posterior contrasts comparing Groups 2–5 with Group 1 were calculated separately within the synch and asynch conditions.

#### Distribution and Transitions of Thought Groups

To examine whether experimental condition was associated with differences in the overall distribution and temporal organization of thought groups, we performed Bayesian analyses of thought-group report distributions and transitions between consecutive thought probes. All five thought groups were included in these analyses.

- Distribution: Condition was included as the fixed effect to test whether the relative distribution of reports across the five thought groups differed between the synch and asynch conditions.
- Transitions: Current thought group, condition, and their interaction were included as fixed effects, with the Subsequent thought group as the outcome. Transitions were defined between consecutive thought probes within each participant and condition. Transition matrices were additionally constructed for each condition to visualize descriptive transition patterns.

#### Model Specifications and Prior Distributions

Appropriate probability distributions and link functions were selected based on the characteristics of each outcome. Gaussian distributions with identity links were used for the continuous probe ratings, modeled response-option probabilities in the categorical probes, RTCV, and standardized RR interval. Response errors were modeled using a binomial distribution with a logit link, with the number of errors specified relative to the total number of tone events within each trial. Thought-group report counts were modeled using a multinomial distribution with a logit link, and transitions were modeled using a categorical distribution with a logit link.

Participant was included as a random intercept in all models. Trial was additionally included as a random intercept in the continuous phenomenological-probe models, whereas participant-specific random slopes for Condition were included in the RTCV, response-error, and standardized-RR models.

For the Gaussian phenomenological-probe models, Normal (0, 2.5) priors were specified for population-level coefficients. Default *brms* priors were retained for the multinomial, categorical, RTCV, and response-error models. For the standardized-RR model, Normal (0, 1) priors were specified for population-level coefficients, Student-*t*(3, 0, 1) priors for the intercept, group-level standard deviations, and residual standard deviation, and an LKJ(1) prior for random-effects correlations.

#### Computation and Inference

Posterior distributions were estimated using Markov Chain Monte Carlo (MCMC) sampling with four chains. For the analyses of subjective thought characteristics, thought-group distributions, and transitions, each chain consisted of 2,000 iterations, with the first 1,000 serving as warm-up. For the objective behavioral and cardiac analyses, each chain consisted of 4,000 iterations, with the first 1,000 serving as warm-up, yielding 12,000 post-warm-up draws per model. Convergence was assessed using the R-hat statistic, with values of R-hat ≤ 1.01 considered acceptable.

The target acceptance rate (*adapt_delta*) was set to 0.99, and the maximum tree depth (*max_treedepth*) was set to 15. Effects were interpreted as supported when the 95% credible interval did not include zero.

#### Bayesian Multinomial Model for Thought Report Distribution and HCT Performance

To investigate whether the distribution of thought reports varied as a function of experimental condition and cardiac interoceptive accuracy, we fitted a Bayesian multinomial mixed-effects model to the vector of report counts assigned to the five thought groups within each participant and condition. This model included all five thought groups: On-task thought (Group 1), Stimulus-dependent thought (Group 2), Self-related thought (Group 3), Task/Self-unrelated thought (Group 4), and Thought absence (Group 5).

For each participant and condition, the dependent variable was the vector of report counts assigned to the five thought groups. The total number of valid thought probes within that participant and condition was included as the number of trials in the multinomial model. Condition, standardized HCT performance, and their interaction were included as fixed effects. Participant was included as a random intercept to account for individual differences in overall report distribution. HCT scores were z-scored before analysis to improve model convergence and facilitate the interpretation of interaction terms. When a participant made no reports of a given thought group in a given condition, the corresponding count was coded as 0.

A multinomial distribution with a logit link function was used. The model was fitted using brms with the default priors implemented for multinomial models, including flat priors for population-level coefficients and Student-*t* (3, 0, 2.5) priors for category-specific intercepts and group-level standard deviations. Posterior distributions were estimated using MCMC methods with four chains, each consisting of 4,000 iterations, including a 1,000-iteration warm-up period. The target acceptance rate (adapt_delta) was set to 0.99, and the maximum tree depth (max_treedepth) was set to 15. Convergence was assessed using the R-hat statistic; values of R-hat ≤ 1.01 were considered indicative of acceptable convergence. Statistical inference was based on 95% credible intervals, and effects were interpreted as supported when the 95% credible interval did not include zero.

Posterior expected values were summarized as predicted report counts per 15 thought probes and as predicted report probabilities. To characterize the Condition × HCT effect for each thought group, we calculated the difference in the synch–asynch contrast between high and low HCT performance, defined as HCT z-scores of +1 and −1, respectively. The Condition × HCT contrast was therefore defined as [(Synch − Asynch) at high HCT] − [(Synch − Asynch) at low HCT]. For each contrast, the posterior mean and 95% CI were summarized from the posterior draws. We additionally calculated the probability of direction, P(direction), as the proportion of posterior draws having the same sign as the posterior mean contrast.

#### Calculation of Heartbeat-Evoked Potentials (HEP)

To examine heartbeat-related cortical activity across thought groups and experimental conditions, heartbeat-evoked potentials (HEPs) were calculated from EEG epochs time-locked to ECG R peaks. EEG data were epoched from −100 to 500 ms relative to each R peak. To minimize contamination of the baseline by cardiac electrical activity surrounding the R wave, the mean amplitude from −100 to −50 ms was subtracted from each epoch.

The analysis focused on heartbeat-related cortical activity during ongoing thought episodes rather than phasic responses immediately following sound omissions. R-peak-locked epochs immediately following sound omissions, which were analyzed separately in the physiological-validation analysis (Sakuragi et al., 2026), were therefore excluded. Intervals from thought-probe onset until completion of the probe response were also excluded to minimize contamination by probe-related cognitive and motor activity. The remaining R-peak-locked epochs were assigned to the thought group reported at the subsequent thought probe.

Because tones in the synch condition were presented at a fixed latency of 200 ms after the R peak, sound-evoked activity was systematically time-locked to the R peak in this condition. We therefore estimated the auditory-evoked contribution using the asynch condition, in which sound onset was temporally independent of ongoing R peaks. For each participant, all available tone-present AEP epochs from the asynch condition were pooled across the five thought groups and averaged to obtain a participant-specific AEP template. The template was aligned to the R-peak-locked HEP time axis such that AEP onset corresponded to 200 ms after the R peak and was subtracted from synch HEP epochs from sound onset onward. No additional AEP subtraction was applied to the asynch condition because auditory events were not systematically time-locked to the R peaks and were therefore expected to average out in the R-peak-locked waveform.

The region of interest (ROI) and analysis window were defined using a functional localizer that was independent of the thought group and condition effects tested in the subsequent analyses. The localizer was conducted separately for the synch and asynch tone-present conditions using retained non-omission R-peak-locked epochs. Pooled-AEP-corrected waveforms were used for the synch condition, whereas the asynch R-peak-locked waveforms were used without additional subtraction. Inspection of the estimated auditory contribution indicated that sound-evoked activity became substantially more pronounced after approximately 300 ms following the R peak; the localizer search was therefore restricted to activity up to 300 ms to minimize residual auditory contamination. The contribution of auditory activity across the full waveform is shown in the Supplementary Material.

Within this temporal range, R-peak-locked EEG amplitudes were tested for negative deviations from 0 μV using nonparametric cluster-based permutation tests implemented in the FieldTrip toolbox for MATLAB (Maris & Oostenveld, 2007; Oostenveld et al., 2011). Samples exceeding a cluster-forming threshold of *p* < .05 were clustered across neighboring electrodes located within 40 mm, and cluster-level significance was evaluated using 2,000 Monte Carlo permutations with a one-tailed cluster-level threshold of α = .05. The final ROI and analysis window were defined as the spatiotemporal region shared by significant negative clusters identified in both conditions.

The functional localizer identified a right frontal-to-parietal cluster spanning 200–300 ms after the R peak, comprising electrodes E1, E2, E3, E4, E5, E6, E7, E8, E9, and E11. Repeating the localizer after applying thought-group-specific AEP correction to the synch data yielded the same ROI and time window. For each retained heartbeat epoch, HEP amplitude was calculated as the mean EEG amplitude across this ROI and the 200–300 ms interval. Cell-level HEP amplitudes were then calculated separately for each available participant × condition × thought-group combination. All five thought groups were retained in the HEP analyses. The number of retained R-peak-locked epochs varied across participants, thought groups, and conditions. Table 2 presents the mean, standard deviation, minimum, and maximum number of retained epochs for each thought group in the synch and asynch conditions.

**Table 2.**
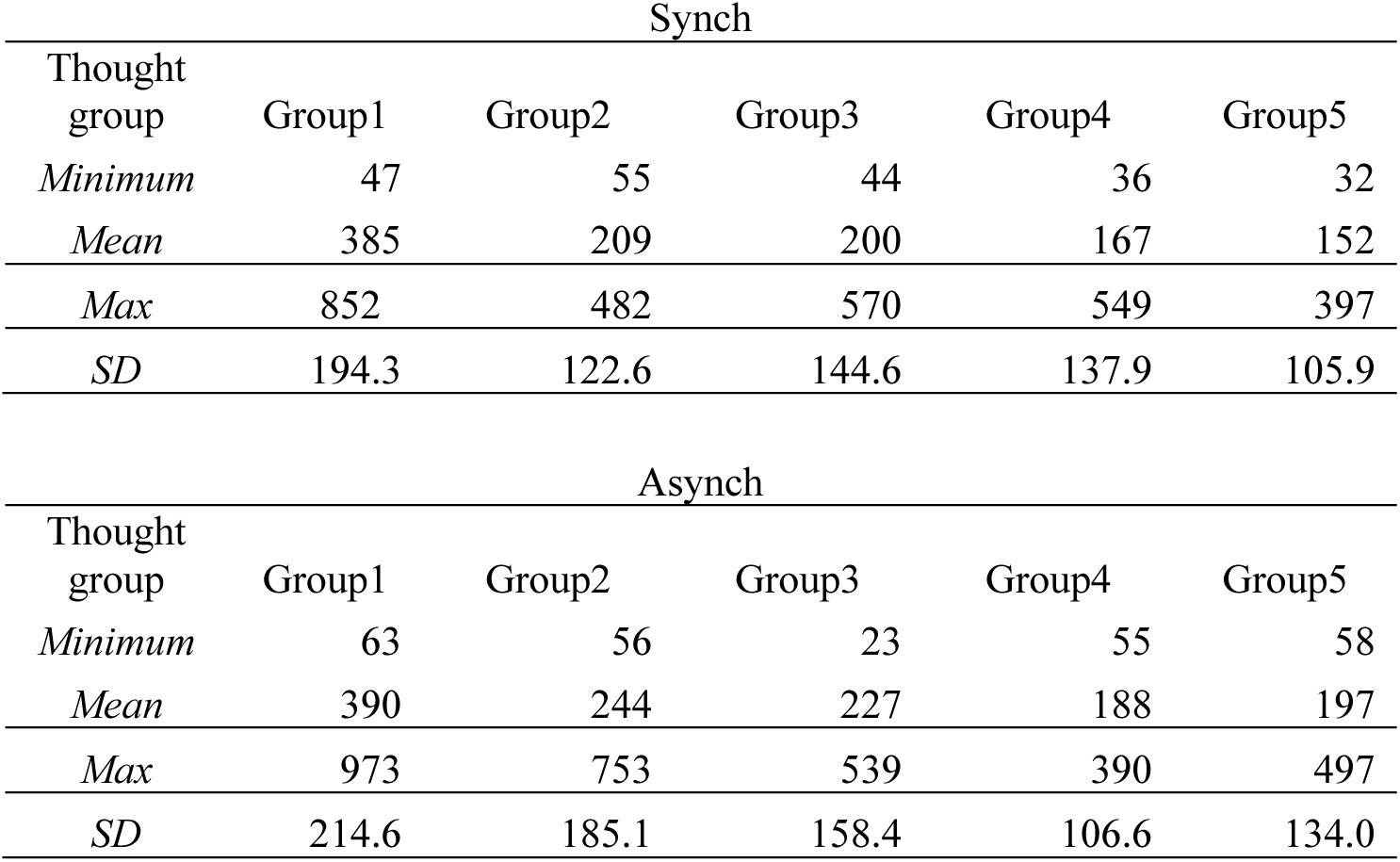
Mean number of R-peak-locked epochs retained for HEP averaging per participant by condition and thought group.

#### Statistical Analysis of HEP Amplitude Primary HEP model and sampling uncertainty

To examine whether HEP amplitude varied across thought groups and experimental conditions, a Bayesian linear mixed-effects model was specified as the primary HEP analysis using the brms package in R. The dependent variable was the mean HEP amplitude within the predefined ROI and 200–300 ms time window for each participant, condition, and thought group. All five thought groups were included in the analyses, with Group 1 (On-task) and the asynch condition specified as the reference levels.

The number of heartbeat epochs contributing to each mean HEP estimate varied across participants, conditions, and thought groups because of differences in thought-report frequency and artifact rejection. Moreover, the precision of an averaged HEP depends not only on the number of contributing epochs but also on the variability of HEP amplitude across those epochs. We therefore quantified the sampling uncertainty of each mean HEP estimate. First, for each retained heartbeat epoch, HEP amplitude was calculated by averaging EEG activity across the predefined ROI and the 200–300 ms time window. For each participant, condition, and thought group, these epoch-level HEP amplitudes were then averaged to obtain the HEP value entered into the model. The standard error of this mean was calculated as the standard deviation of the epoch-level HEP amplitudes divided by the square root of the number of retained epochs (SE=SD/√*n*). Thus, mean HEP estimates based on fewer or more variable epochs were associated with larger standard errors, whereas estimates based on more numerous and less variable epochs were associated with smaller standard errors.

These estimate-specific standard errors were incorporated into the Bayesian model using the se() response specification in brms (Bürkner, 2017), with an additional residual variance term estimated (sigma = TRUE). This allowed differences in sampling uncertainty among the mean HEP estimates to be considered while retaining an additional residual variance term. The standardized number of retained epochs was also included as a covariate. This term served a different purpose from the standard-error specification: whereas the standard error represented uncertainty in the estimated mean HEP amplitude, the epoch-count covariate accounted for any systematic association between the number of contributing epochs and HEP amplitude itself.

The model included thought group, condition, and their interaction as fixed effects, together with standardized epoch count as a covariate. Participant was included with a random intercept to account for repeated observations within participants and a random slope for Condition to allow the condition effect to vary across individuals. The model was specified as:

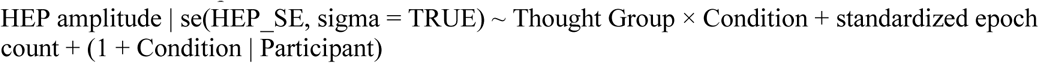

#### Sensitivity analyses

Sensitivity analyses were conducted to assess the robustness of the HEP results to two analytic choices: the method used to correct for auditory-evoked activity and the explicit modelling of sampling uncertainty in the mean HEP estimates.

First, to assess whether the results depended on the AEP-correction procedure, the primary HEP model was repeated using three versions of the synch HEP data: uncorrected HEPs, HEPs corrected using participant-specific pooled AEP templates, and HEPs corrected using participant-by-thought-group-specific AEP templates. To ensure that differences between these analyses reflected the correction procedure rather than differences in data availability, all three models were fitted to the same set of participant × thought-group observations for which both experimental conditions and all three correction variants were available.

Participant-specific pooled AEP correction was used in the primary analysis because pooling all available asynch tone-present epochs across thought groups provided a more stable estimate of each participant’s auditory response. Thought-group-specific correction was examined as an alternative that allowed the auditory response to vary across thought groups, and the uncorrected analysis was used to assess whether the conclusions depended on AEP subtraction itself.

Second, to assess whether the results depended on explicitly modelling differences in the sampling uncertainty of the mean HEP estimates, the primary model was repeated without the estimate-specific standard-error term while retaining the same observations, predictors, and random-effects structure.

For all HEP models, Normal(0, 10) priors were assigned to the non-intercept population-level coefficients, and half-Cauchy(0, 2) priors were assigned to the standard deviations of the participant-level random effects. The brms default priors were retained for the intercept, residual standard deviation, and random-effects correlation structure: Student-t(3, −1, 2.5) for the intercept, Student-t(3, 0, 2.5) for the residual standard deviation, and LKJ(1) for the random-effects correlation matrix. Posterior distributions were estimated using four Markov chain Monte Carlo chains, each consisting of 4,000 iterations, including 1,000 warm-up iterations. The target acceptance probability was set to .95. Model convergence and sampling diagnostics were assessed using the R-hat statistic and the presence of divergent transitions. R-hat values below 1.01 were considered acceptable.

Statistical inference was based on 95% CI; effects were considered supported when the 95% CI did not include zero.

## Results

### Report proportions of each thought category

Descriptive statistics for the report proportions of each thought category are shown separately for the synch and asynch conditions in Tables 3 and 4. In both conditions, reports were most frequently assigned to task-focused, task-related, self-related, and interoceptive categories, whereas exteroceptive reports and forgetfulness were infrequent. The overall descriptive pattern was broadly similar across the asynch and synch conditions, with task-focused reports being slightly more frequent in the synch condition and task-related reports being slightly more frequent in the asynch condition. Considerable between-participant variability was observed across categories, as indicated by the wide ranges and standard deviations.

**Table 3.**
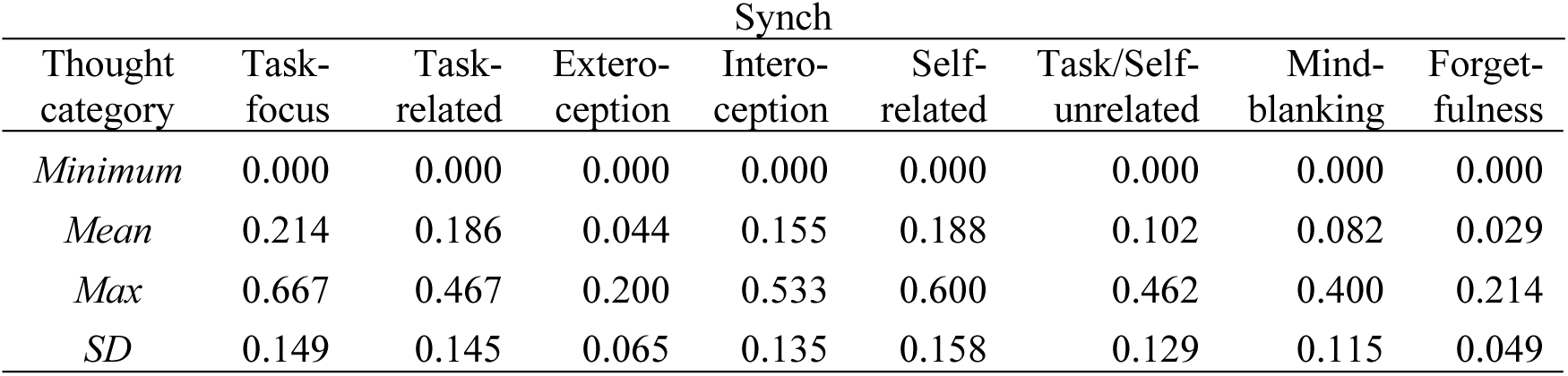
Report proportions of each thought category in the synch condition.

| Thought category | Synch |  |  |  |  |  |  |  |
| --- | --- | --- | --- | --- | --- | --- | --- | --- |
|  | Task-focus | Task-related | Exteroception | Interoception | Self-related | Task/Self-unrelated | Mind-blanking | Forgetfulness |
| *Minimum* | 0.000 | 0.000 | 0.000 | 0.000 | 0.000 | 0.000 | 0.000 | 0.000 |
| *Mean* | 0.214 | 0.186 | 0.044 | 0.155 | 0.188 | 0.102 | 0.082 | 0.029 |
| *Max* | 0.667 | 0.467 | 0.200 | 0.533 | 0.600 | 0.462 | 0.400 | 0.214 |
| *SD* | 0.149 | 0.145 | 0.065 | 0.135 | 0.158 | 0.129 | 0.115 | 0.049 |

**Table 4.** Report proportions of each thought category in the asynch condition.

| Thought category | Asynch |  |  |  |  |  |  |  |
| --- | --- | --- | --- | --- | --- | --- | --- | --- |
|  | Task-focus | Task-related | Exteroception | Interoception | Self-related | Task/Self-unrelated | Mind-blanking | Forgetfulness |
| *Minimum* | 0.000 | 0.000 | 0.000 | 0.000 | 0.000 | 0.000 | 0.000 | 0.000 |
| *Mean* | 0.191 | 0.196 | 0.038 | 0.144 | 0.183 | 0.109 | 0.108 | 0.030 |
| *Max* | 0.733 | 0.467 | 0.200 | 0.467 | 0.533 | 0.357 | 0.467 | 0.200 |
| *SD* | 0.202 | 0.131 | 0.062 | 0.130 | 0.148 | 0.114 | 0.136 | 0.058 |

### Characteristics of thought groups across experimental conditions

We first examined whether the four thought groups for which phenomenological ratings were available showed distinct profiles on the continuous thought probes (Figure 4): Group 1 (On-task), Group 2 (Stimulus-dependent), Group 3 (Self-related), and Group 4 (Task/Self-unrelated). Bayesian linear mixed-effects models were fitted separately for each condition and probe, with Group 1 serving as the reference category; full model estimates are reported in Tables A1–A10 (Supplementary). Corresponding analyses of the categorical thought probes are reported in Figure A1 and Tables A11–A18 (Supplementary).

**Figure 4.**
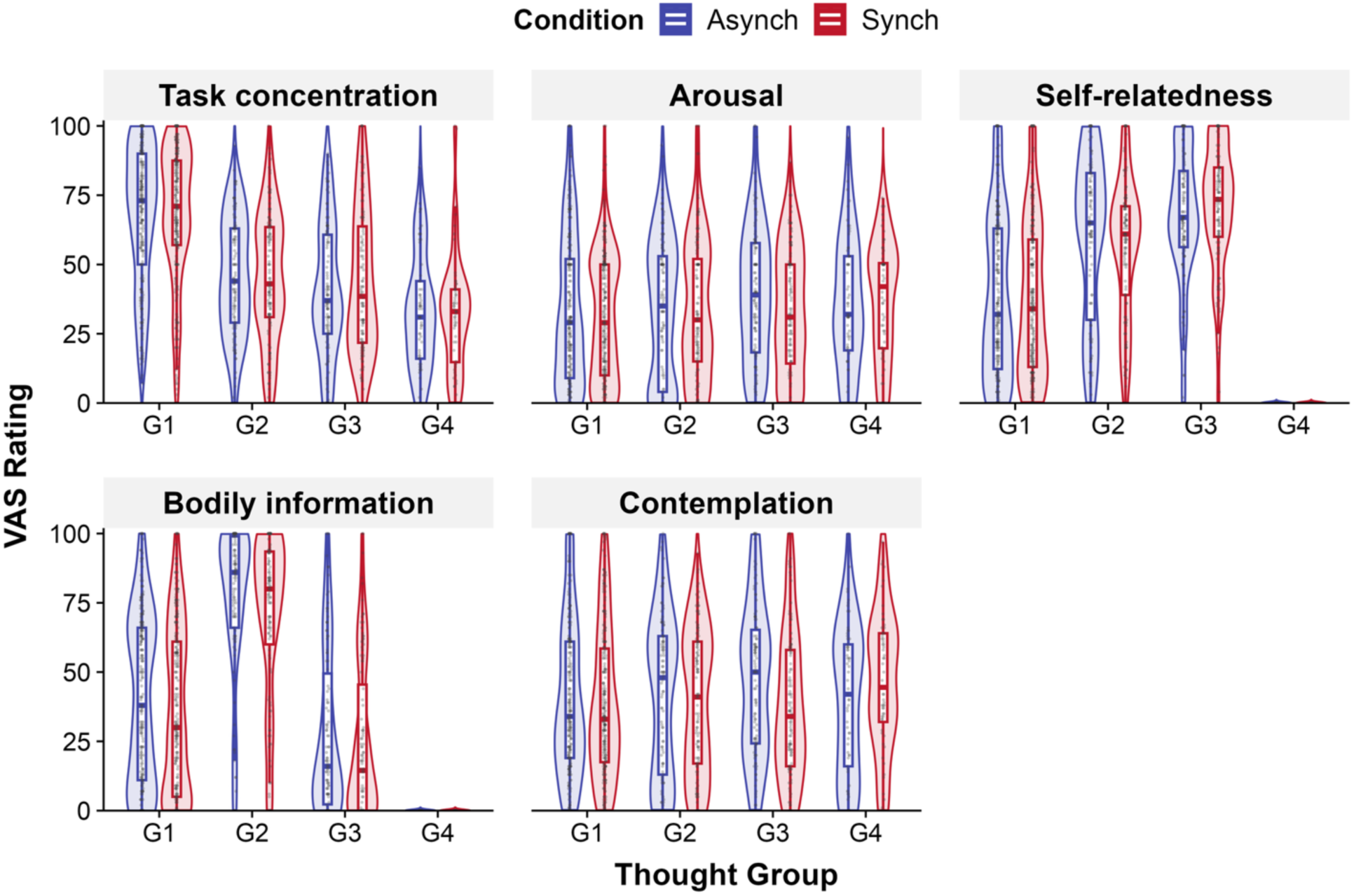
VAS ratings of continuous probes for each thought group across experimental conditions. Violin plots, box plots, and individual data points show VAS ratings for task concentration, arousal, self-relatedness, bodily information, and contemplation in the asynch and synch conditions. Within each thought group, the asynch and synch conditions are shown side by side. G1 = On-task; G2 = Stimulus-dependent thought; G3 = Self-related thought; G4 = Task/Self-unrelated thought. Bayesian linear mixed-effects models were used to examine group differences within each condition, with On-task (G1) as the reference category. Across conditions, G1 showed higher task concentration, G2 showed higher bodily information ratings, and G3 showed higher self-relatedness and contemplation. G2 and G3 also showed lower task concentration and higher self-relatedness than G1. In the synch condition, G2 and G3 showed higher arousal and contemplation relative to G1. Overall, these patterns support the interpretation that the predefined thought groups captured distinct subjective profiles. Alt text. Five-panel violin plots compare VAS ratings across four thought groups in the asynch and synch conditions for task concentration, arousal, self-relatedness, bodily information, and contemplation. Group 1 shows the highest task concentration, Group 2 shows the highest bodily-information ratings, and Group 3 shows high self-relatedness and contemplation. Group 4 shows very low self-relatedness and bodily-information ratings. The overall pattern of differences among thought groups is broadly similar across the asynch and synch conditions, with some condition-related variation in arousal and contemplation.

Across both experimental conditions, the continuous-probe profiles were consistent with the intended definitions of the thought groups. Group 1 was characterized by the highest level of task concentration. Group 2 was primarily composed of interoceptive reports and showed higher bodily-information ratings, consistent with a body-oriented form of stimulus-dependent thought. Group 3 showed high self-relatedness and contemplation, consistent with self-related thought. Group 4 showed low self-relatedness and task concentration, consistent with task- and self-unrelated thought. These profiles indicate that the predefined thought groups captured qualitatively distinct forms of ongoing mental experience.

The overall structure of group differences was broadly similar between the asynch and synch conditions. Although within-condition comparisons against Group 1 indicated that Groups 2 and 3 showed higher arousal and contemplation in the synch condition, the basic interpretation of each thought group was preserved across conditions. Supplementary analyses of categorical probes showed broadly consistent qualitative differentiation among groups (Figure A1). These results support the qualitative differentiation of Groups 1–4 used in the subsequent analyses.

Objective behavioral and cardiac measures provided additional characterization of the five thought groups (Supplementary Figure A2; Tables A19–A23). Response errors were infrequent overall, but posterior contrasts indicated lower error probabilities in the synch than in the asynch condition for Groups 1–4, with a similar directional difference in Group 5 whose 95% credible interval included zero (Table A21). Standardized RR intervals were lower in Groups 2–5 than in Group 1 in the asynch condition, whereas in the synch condition this contrast was clearly supported only for Group 3; however, there was no clear evidence for Condition × Thought Group interactions in RR interval (Tables A22–A23). Together, these patterns suggest that the synch condition was generally associated with fewer response errors, while Self-related thought was accompanied by relatively shorter RR intervals than On-task thought across conditions. Full results, including RTCV, are reported in the Supplementary Material.

### Report frequencies and transition patterns of thought groups

We next examined whether the synch condition altered the distribution and transition patterns of all five thought groups, including Group 5 (Thought absence). Group-level report proportions are shown in Figure 5, and full model estimates for the report-distribution analysis are provided in Table A24 (Supplementary). In both conditions, On-task thought (Group 1) was the most frequently reported thought group. A Bayesian multinomial mixed-effects model did not provide strong evidence for condition-related differences in the relative probability of any thought group, as all 95% CIs for the condition effects included zero. Thus, the overall distribution of reports assigned to the five thought groups was broadly similar across conditions.

**Figure 5.**
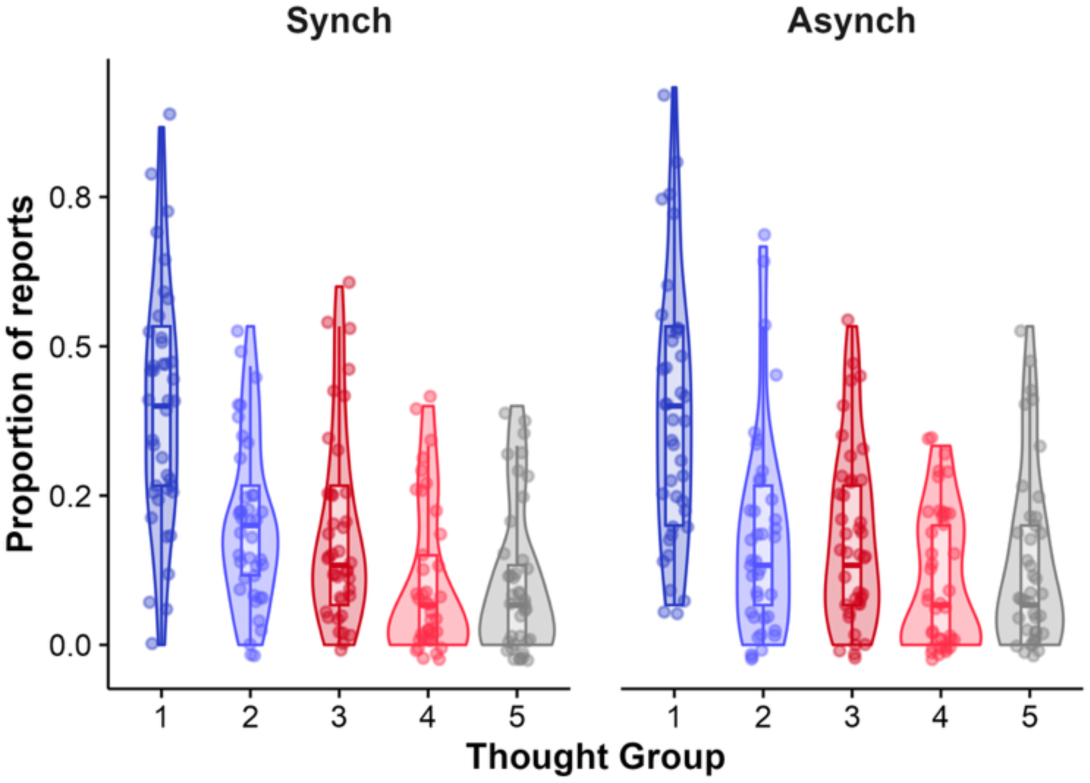
Report proportions of each thought group by experimental condition.. The x-axis represents the five thought groups, and the y-axis represents the proportion of reports for each participant. Violin plots, box plots, and individual data points show the distributions of report proportions in the synch and asynch conditions. On-task thought (Group 1) showed the highest report proportion in both conditions. The overall distributions were broadly similar across conditions, although descriptive differences were observed in the relative proportions of some thought groups. Alt text: Two-panel violin plots show the distribution of report proportions for five thought groups in the synch and asynch conditions. On-task thought (Group 1) has the highest report proportion in both conditions. Stimulus-dependent thought (Group 2) and Self-related thought (Group 3) occur at intermediate levels, whereas Task/Self-unrelated thought (Group 4) and Thought absence (Group 5) are generally less frequent. The overall distribution pattern is broadly similar across conditions, with substantial between-participant variability within each thought group.

We then examined transition patterns between thought groups (Figure 6; Table A25). Figure 6 provides a descriptive visualization of transition probabilities in each condition, whereas Table A25 (Supplementary) summarizes synch–asynch contrasts derived from the Bayesian categorical mixed-effects model. None of the transition-specific posterior contrasts provided conclusive evidence for a condition effect, as all 95% CIs included zero. Nevertheless, the largest directional contrasts suggested a pattern centered on Stimulus-dependent thought (Group 2). In the synch condition, transitions from On-task thought to Stimulus-dependent thought, from Stimulus-dependent thought to Self-related thought, and from Stimulus-dependent thought back to On-task thought tended to be more likely than in the asynch condition. Conversely, transitions from Stimulus-dependent thought and Self-related thought to Thought absence tended to be less likely in the synch condition.

**Figure 6.**
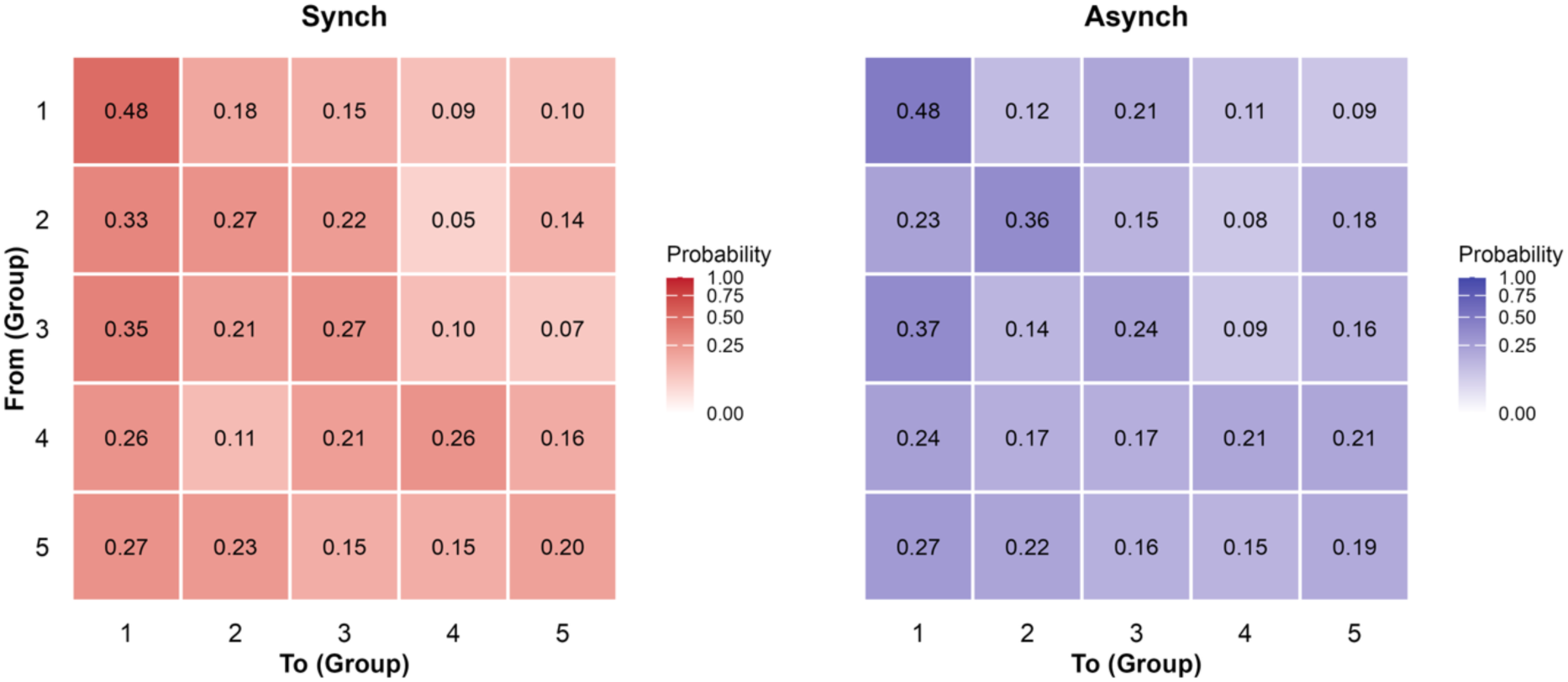
Thought-group transition matrices by experimental condition.. Rows indicate the source thought group, and columns indicate the destination thought group. Numerical values within each tile represent the observed transition probability from the source group to the destination group, and color intensity reflects the magnitude of that probability. Descriptively, the synch condition showed relatively more prominent transitions from On-task thought (Group 1) to Stimulus-dependent thought (Group 2), from Stimulus-dependent thought to Self-related thought (Group 3), and from Stimulus-dependent thought back to On-task thought than the asynch condition. Alt text: Two 5 × 5 transition matrices compare observed probabilities of moving between five thought groups in the synch and asynch conditions. Rows indicate the current thought group and columns the subsequent thought group. Self-transitions from On-task thought (Group 1) are the most common in both conditions, with a probability of 0.48. Relative to the asynch condition, the synch condition shows numerically higher transition probabilities between On-task and Stimulus-dependent thought and between Stimulus-dependent and Self-related thought, while transitions from Stimulus-dependent and Self-related thought to Thought absence are numerically lower.

### Relationship between thought group reporting frequency and cardiac interoceptive accuracy

To examine whether cardiac interoceptive accuracy modulated condition-related changes in the distribution of reports assigned to the thought groups, we fitted a Bayesian multinomial mixed-effects model to the vector of report counts assigned to the five thought groups within each participant and condition. Cardiac interoceptive accuracy was indexed by standardized HCT performance. Fixed-effect estimates from the model are summarized in Table A26. Because multinomial logit coefficients are expressed relative to the reference outcome category, we based the main interpretation on posterior predicted Condition × HCT contrasts in report counts and probabilities, summarized in Table 5.

**Table 5.** Posterior predicted Condition × HCT contrasts in thought report distribution.

| Group | Condition × HCT | Condition × HCT, probability | P(direction),<br>probability |
| --- | --- | --- | --- |
| Group 1 | 0.47 [-1.44, 2.40] | 0.031 [-0.096, 0.160] | 0.684 |
| Group 2 | -0.16 [-1.60, 1.27] | -0.011 [-0.107, 0.085] | 0.584 |
| Group 3 | 1.54 [0.08, 3.09] | 0.103 [0.005, 0.206] | 0.981 |
| Group 4 | -1.59 [-2.84, -0.63] | -0.106 [-0.189, -0.042] | >.999 |
| Group 5 | -0.27 [-1.33, 0.74] | -0.018 [-0.088, 0.049] | 0.701 |

*Note.* Condition × HCT contrasts were defined as [(Synch – Asynch) at HCT $z = +1$ ] – [(Synch – Asynch) at HCT $z = -1$ ]. Positive values indicate a more positive synch–asynch difference at higher HCT performance. P(direction) represents the posterior probability in the direction of the estimated contrast.

The posterior predicted contrasts indicated that condition-related changes in the distribution of thought reports varied with HCT performance. For Self-related thought (Group 3), the Condition × HCT contrast was positive on both the predicted-count scale (1.54 reports per 15 probes, 95% CI [0.08, 3.09]) and the probability scale (0.103, 95% CI [0.005, 0.206]; P(direction) = .981). For Task/Self-unrelated thought (Group 4), the corresponding contrast was negative (predicted count: −1.59, 95% CI [−2.84, −0.63]; probability: −0.106, 95% CI [−0.189, −0.042]; P(direction) > .999). The contrasts for On-task thought (Group 1), Stimulus-dependent thought (Group 2), and Thought absence (Group 5) had 95% CIs that included zero.

Thus, participants with higher HCT performance showed a greater synch-related increase in Self-related thought reports and a greater synch-related decrease in Task/Self-unrelated thought reports. This pattern indicates that condition-related changes in the distribution of thought reports varied with HCT performance, with higher HCT performance associated with a more positive synch–asynch difference in Self-related thought and a more negative synch–asynch difference in Task/Self-unrelated thought. The relationship between HCT performance and posterior predicted report counts for each thought group is shown in Figure 7.

**Figure 7.**
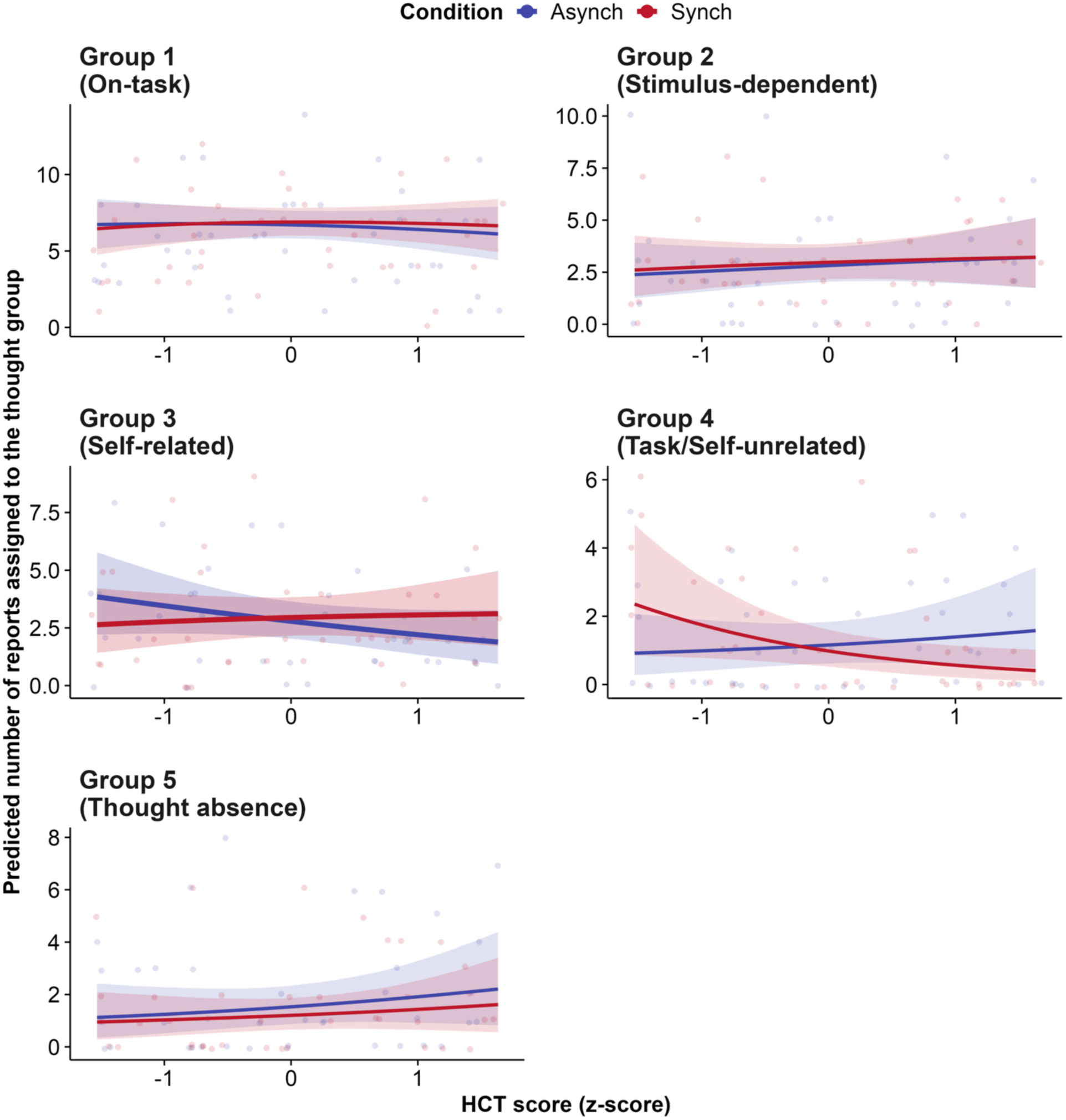
Relationship between cardiac interoceptive accuracy and the predicted number of reports assigned to each thought group. Each panel shows one thought group included in the Bayesian multinomial mixed-effects model: On-task thought (Group 1), Stimulus-dependent thought (Group 2), Self-related thought (Group 3), Task/Self-unrelated thought (Group 4), and Thought absence (Group 5). The x-axis represents cardiac interoceptive accuracy, operationalized as standardized HCT performance. The y-axis represents the posterior predicted number of reports assigned to each thought group. Individual data points show observed report counts for each participant and condition and are jittered for visibility. Solid lines indicate posterior mean predictions, and shaded regions indicate 95% CI. Y-axis ranges vary across panels to improve visualization of within-group associations. The model showed a positive Condition × HCT contrast for Self-related thought and a negative Condition × HCT contrast for Task/Self-unrelated thought, indicating a condition- and HCT-dependent redistribution of thought reports. Alt text: Five-panel plots show posterior-predicted report counts for each thought group as a function of standardized HCT score in the synch and asynch conditions. Condition-related differences vary most clearly for Self-related thought (Group 3) and Task/Self-unrelated thought (Group 4): with increasing HCT score, predicted Self-related thought becomes relatively more frequent in the synch than in the asynch condition, whereas predicted Task/Self-unrelated thought becomes relatively less frequent. Associations are weaker for On-task thought, Stimulus-dependent thought, and Thought absence. Shaded bands indicate uncertainty around the posterior mean predictions, and individual points show observed report counts.

### Neural activity associated with each thought group

To examine whether HEP amplitude varied across thought groups and heartbeat-synchronization conditions, we fitted the Bayesian linear mixed-effects model described above to mean HEP amplitude within the predefined ROI and 200–300 ms interval. The model accounted for the sampling uncertainty of each mean HEP estimate, included standardized epoch count as a covariate, and allowed the condition effect to vary across participants. Grand-average topographies and HEP waveforms are shown in Figures 8A and 8B, respectively, and posterior Synch–Asynch contrasts and model-estimated HEP amplitudes are shown in Figures 8C and 8D. Key posterior contrasts are summarized in Table 6, and the full model coefficients are reported in Table A27 (Supplementary).

**Figure 8.**
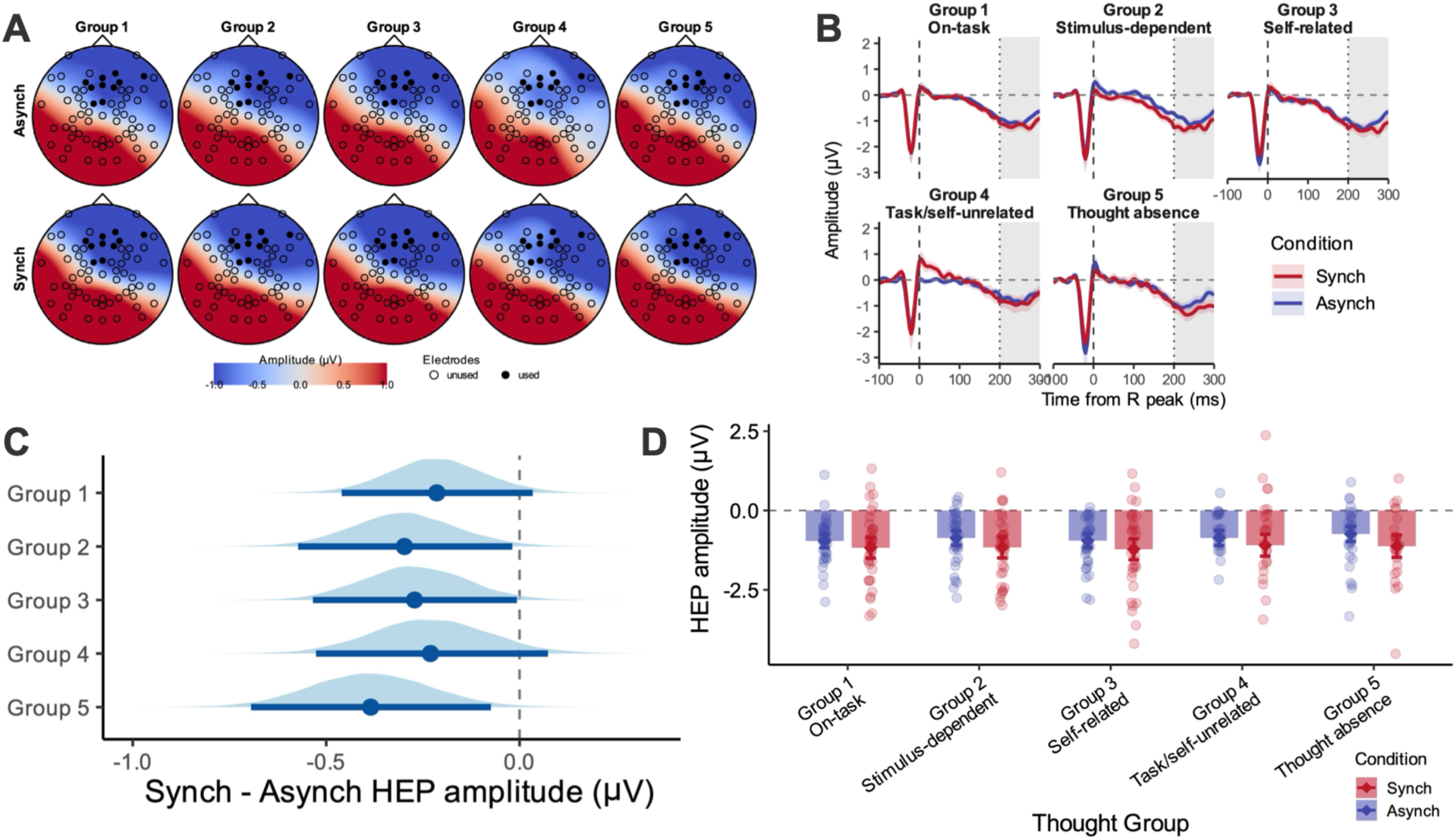
HEP amplitude by thought group and heartbeat-synchronization condition. (A) Grand-average topographic maps of HEP amplitude within the 200–300 ms window for each thought group and condition. Filled electrodes indicate the predefined ROI used for HEP quantification. (B) Grand-average HEP waveforms for each thought group and condition. The grey shaded area indicates the 200–300 ms analysis window. (C) Posterior distributions of the within-group condition contrasts in HEP amplitude (Synch − Asynch), shown separately for each thought group. Points indicate posterior means, and horizontal lines indicate 95% credible intervals. (D) Posterior-predicted HEP amplitudes within the 200–300 ms window for each thought group and condition. Points indicate individual participant estimates, diamonds indicate posterior means, and error bars indicate 95% credible intervals. Alt text: Four-panel figure summarizes heartbeat-evoked potentials (HEPs) across five thought groups in the synch and asynch conditions. Panel A shows scalp topographies of mean HEP amplitude from 200–300 ms after the R peak, with the predefined right frontal-to-parietal ROI marked by filled electrodes; broadly similar spatial patterns are observed across thought groups and conditions. Panel B shows grand-average HEP waveforms, with synch and asynch waveforms following similar time courses and the synch condition tending toward greater negativity within the shaded 200–300 ms analysis window. Panel C shows posterior distributions of the Synch–Asynch HEP contrast for each thought group; all estimates are negative, with 95% credible intervals excluding zero for Stimulus-dependent thought, Self-related thought, and Thought absence, but including zero for On-task and Task/Self-unrelated thought. Panel D shows individual and posterior-predicted HEP amplitudes for each condition and thought group, illustrating the generally more negative HEP amplitudes in the synch condition.

**Table 6.** Posterior contrasts of HEP amplitude between the synch and asynch conditions across thought groups.

| Thought group | Synch – Asynch |  | Difference in condition |  |
| --- | --- | --- | --- | --- |
| | contrast, Estimate ($\mu\text{V}$) | 95% CI | effect contrast vs. Group 1, Estimate ($\mu\text{V}$) | 95% CI |
| Group 1 | -0.214 | [-0.460, 0.034] |  |  |
| Group 2 | -0.298 | [-0.573, -0.019] | -0.084 | [-0.288, 0.120] |
| Group 3 | -0.272 | [-0.534, -0.007] | -0.058 | [-0.247, 0.130] |
| Group 4 | -0.230 | [-0.526, 0.073] | -0.016 | [-0.254, 0.222] |
| Group 5 | -0.385 | [-0.694, -0.074] | -0.171 | [-0.419, 0.081] |

The Group × Condition interactions did not provide clear evidence that the effect of heartbeat-synchronization condition differed across thought groups; the 95% CIs for all corresponding contrasts included zero (Table 6). At the reference level of the asynch condition, Thought absence (Group 5) showed a more positive HEP amplitude than On-task thought (Group 1), whereas differences between Group 1 and the other thought groups were not clearly supported (Table A27).

Within-group posterior contrasts nevertheless showed a consistent shift toward more negative HEP amplitudes in the synch condition across all five thought groups (Figure 8C; Table 6). For Stimulus-dependent thought (Group 2), Self-related thought (Group 3), and Thought absence (Group 5), the 95% CI excluded zero, providing evidence that HEP amplitudes were more negative in the synch than in the asynch condition within these thought groups. The estimated contrasts for On-task thought (Group 1) and Task/self-unrelated thought (Group 4) were in the same negative direction but were estimated with greater uncertainty and included zero. Thus, the synch condition was associated with a supported negative HEP shift in Groups 2, 3, and 5, together with a similar directional tendency in Groups 1 and 4. However, because the Group × Condition interactions were not supported, these findings do not indicate that the condition effect was selectively stronger in Groups 2, 3, or 5 than in the other thought groups.

The conclusion regarding thought-group specificity was consistent across the principal sensitivity analyses. In the matched-sample comparison of AEP-correction procedures, Synch–Asynch contrasts were negative across all five thought groups for uncorrected, pooled-AEP-corrected, and thought-group-specific AEP-corrected HEPs, although the magnitude and interval support of the within-group contrasts varied across correction procedures (Supplementary Figure A3; Table A28). Importantly, none of the correction procedures provided clear evidence that the condition effect differed across thought groups (Supplementary Table A28). Similarly, when estimate-specific sampling uncertainty was not explicitly modelled, within-group contrasts remained negative but their 95% credible intervals included zero, while no between-group difference in the condition effect was supported (Supplementary Table A29). Thus, the evidence for the magnitude of the overall synch-related negative shift was sensitive to analytic specification, whereas the absence of robust thought-group-specific modulation was consistent across analyses.

## Discussion

In the present study, we examined whether covert heartbeat-related bodily processing, as a continuously varying state factor operating largely outside explicit awareness, was associated with changes in the organization of spontaneous thought. Participants performed an auditory attention task incorporating a heartbeat-synchronous sound condition (synch), in which heartbeat-synchronous omissions have been shown to modulate cardiac responses and heartbeat-related cortical processing (Sakuragi et al., 2026), and a heartbeat-asynchronous condition (asynch), while ongoing thought was sampled using intermittent thought probes. We examined condition-related differences in thought-state distribution and transition patterns, their variation with cardiac interoceptive accuracy indexed by HCT performance, and HEP amplitude across thought states and conditions. The overall distribution of thought groups was broadly similar across conditions, but directional transition patterns suggested closer temporal links among on-task, body-oriented stimulus-dependent, and self-related thought in the synch condition. Among participants with higher HCT performance, the synch condition was associated with more self-related thought and less task/self-unrelated thought relative to the asynch condition. HEPs showed a synch-related negative shift across several thought groups, with no clear evidence that the magnitude of this condition effect differed among thought groups. Together, these findings suggest that heartbeat-related bodily processing may contribute to the ongoing flow of spontaneous thought, with bodily information becoming more influential in thought transitions and in the distribution of self-related content in individuals with greater cardiac interoceptive accuracy.

The condition-related differences in transition patterns were concentrated around stimulus-dependent thought. In the synch condition, transitions connecting this state with both on-task and self-related thought tended to be more likely than in the asynch condition. Group 2 consisted predominantly of interoceptive reports and was characterized by relatively high bodily-information ratings. This pattern therefore suggests that heartbeat-related modulation may have increased the influence of bodily information on the ongoing flow between task-focused and self-related thought. However, the credible intervals for the individual transition contrasts included zero, so this finding should be interpreted as a directional pattern in the overall organization of thought transitions rather than evidence for a specific transition pathway.

The HCT findings suggest that the influence of heartbeat-related modulation on ongoing thought may depend on individual differences in cardiac interoception. If HCT performance is taken as a behavioral index related to cardiac interoceptive accuracy, the greater relative occurrence of self-related thought and lower occurrence of task/self-unrelated thought in the synch condition among participants with higher HCT performance may indicate that subtle changes in cardiac activity and its neural processing were more strongly reflected in their ongoing thought. This interpretation is consistent with previous studies linking heartbeat-related neural processing to self-related spontaneous thought and bodily self-consciousness (Babo-Rebelo et al., 2016; Park et al., 2018). The subjective and cardiac characteristics of self-related thought further complement this account. These thoughts were characterized by high self-relatedness and contemplation, and in the synch condition they were also associated with relatively higher subjective arousal than on-task thought. Across conditions, they additionally occurred during relatively shorter RR intervals, indicating a tendency to coincide with faster cardiac states within each participant’s task-wide range. Taken together, the self-related thoughts captured here may have reflected a relatively activated and elaborative form of self-focused processing that was accompanied by changes in ongoing cardiac state. These findings do not establish that cardiac changes caused self-related thought, but they support a close association between sensitivity to cardiac signals, ongoing cardiac state, and the emergence of self-related cognition.

The HEP findings suggest that heartbeat-related cortical activity reflected condition-related differences in ongoing cardiac signal processing rather than the self-relatedness of ongoing thought. Although HEP amplitude differed between the synch and asynch conditions across several thought groups, the magnitude of this condition-related difference did not vary reliably with thought group. Thus, the expected selective association between HEP modulation and self-related thought was not supported. This pattern suggests that neural processing of cardiac signals differed between the experimental conditions, while the emergence of self-related thought may have depended on additional factors, including individual cardiac interoceptive accuracy and ongoing cardiac state. HEP therefore provides complementary evidence that heartbeat-related cortical processing differed across the experimental contexts, but it does not identify a neural signature specific to the Self-related thought changes observed here.

In the dynamic framework of spontaneous thought, transitions between mental states are shaped by deliberate and automatic constraints, including processes that operate outside conscious control (Christoff et al., 2016). The present findings extend this framework by suggesting that heartbeat-related bodily processing may constitute one such automatic constraint, biasing both the organization and content of ongoing thought. At the group level, condition-related differences in transition patterns were centered on body-oriented stimulus-dependent thought, whereas across individuals, participants with higher HCT performance showed a stronger shift toward self-related thought in the synch condition. One possible mechanism for these effects is suggested by predictive-processing accounts, in which the impact of interoceptive signals on ongoing processing depends on their context-dependent weighting or precision (Barrett & Simmons, 2015; Seth & Friston, 2016). Although the present study did not directly measure such weighting, individual differences related to cardiac interoception may influence the extent to which heartbeat-related bodily information contributes to ongoing cognition (Ainley et al., 2016).

Several limitations qualify the interpretation of these findings. First, although the present study experimentally manipulated heartbeat-related bodily processing, it does not establish the temporal causal pathway linking individual cardiac responses to specific thought transitions. Thought probes did not provide the precise onset of each transition, and omission-related physiological responses were not linked event by event to subsequent changes in thought state. Moreover, epochs immediately following omissions were excluded from the ongoing RR and HEP analyses, which were intended to characterize cardiac and cortical activity associated with each thought state rather than phasic responses to individual omissions. The present data therefore support condition-level associations between heartbeat-related modulation and thought organization but cannot determine whether a particular cardiac response contributed to a specific transition. Future studies combining higher-temporal-resolution measures of thought-state change with event-level modeling of cardiac, cortical, and thought dynamics will be needed to resolve this causal sequence.

Second, the interpretation of the individual-difference findings is constrained by the use of HCT performance as a behavioral index related to cardiac interoceptive accuracy. Although HCT performance may reflect individual differences in access to cardiac signals, it can also be influenced by processes other than direct perception of ongoing cardiac activity. HEPs provided a complementary cortical index of heartbeat-related processing that did not require explicit heartbeat reports, but neither measure captures the full multidimensional nature of cardiac interoception, including discrimination accuracy, subjective sensibility, and metacognitive correspondence between performance and confidence. Future studies combining complementary behavioral, subjective, and neural measures will be needed to determine which aspects of cardiac interoception are most relevant to the modulation of spontaneous thought.

Third, the precision of thought-group-specific HEP estimates was limited by sparse and uneven numbers of available heartbeat epochs across participant × condition × thought-group cells. Because thought states occurred at different frequencies across participants and conditions, the number of usable epochs contributing to each HEP estimate varied substantially. The primary model accounted for estimate-specific sampling uncertainty and epoch count, and sensitivity analyses consistently provided no clear evidence that the condition effect differed across thought groups, although the magnitude and interval support of the within-group contrasts varied across analytic specifications. Sparse data in some cells may nevertheless have reduced sensitivity to subtle thought-specific differences in HEP modulation. Longer recordings or designs yielding larger and more balanced numbers of epochs for each thought state will be needed to test such effects more reliably.

Fourth, because the heartbeat-related manipulation was embedded within an auditory attention task, subtle differences in auditory or task processing between the synch and asynch conditions cannot be completely excluded as contributors to the observed condition effects. This concern is mitigated by the absence of explicit awareness of the heartbeat–sound relationship and by the limited behavioral evidence for differential task difficulty: although response errors were less frequent in the synch condition, errors were rare overall and RTCV showed no clear condition-related difference. Smaller differences in stimulus processing may nevertheless have contributed to the observed effects. Future studies using alternative timing controls, sensory modalities, and task settings will be important for determining the extent to which these findings generalize beyond the present cardio-auditory paradigm.

## Conclusion

The present study examined whether covert heartbeat-related modulation of cardiac activity and cardiac signal processing was associated with changes in the organization of spontaneous thought. In the condition in which auditory events were synchronized to participants’ heartbeats to induce such modulation, the directional transition pattern was centered on body-oriented Stimulus-dependent thought. Participants with higher HCT performance also showed more Self-related thought and less Task/Self-unrelated thought in this heartbeat-synchronous condition. HEPs likewise differed between these experimental conditions across several thought states, although these differences were not selectively associated with Self-related thought. Taken together, these findings suggest that heartbeat-related bodily signals and their processing may act as an automatic constraint on the ongoing stream of thought, biasing which thought contents and transitions become more likely. The specific physiological and neural pathways through which this influence is implemented remain to be determined.

## CRediT authorship contribution statement

**Mai Sakuragi:** Writing – original draft, Software, Methodology, Data curation, Investigation, Funding acquisition, Formal analysis, Conceptualization.

**Kazushi Shinagawa:** Writing – review & editing, Formal analysis.

**Yuri Terasawa:** Writing – review & editing.

**Yuto Tanaka:** Writing – review & editing, Methodology.

**Satoshi Umeda:** Writing – review & editing, Supervision.

## Ethics approval and consent to participate

The study was approved by the Keio University Research Ethics Committee, Japan (Approval No. 240040000; approved 7 May 2024), and was conducted in accordance with the Declaration of Helsinki and relevant institutional guidelines. All participants provided written informed consent before participation, and participants’ privacy rights were observed.

## Declaration of generative AI and AI-assisted technologies in the manuscript preparation process

During the preparation of this work, the corresponding author used ChatGPT, developed by OpenAI, for language editing, wording refinement, and assistance in improving the clarity and organization of the manuscript. The authors reviewed and edited the manuscript as needed and take full responsibility for its content.

## Acknowledgements

We are grateful to all participants.

## Data availability

The datasets analyzed during the current study are not publicly available because public data sharing was not covered by the approved ethics protocol or by participants’ informed consent. The datasets contain sensitive personal information, including physiological recordings and questionnaire responses, which prevents deposition in a public repository. De-identified data may be made available from the corresponding author upon reasonable request, subject to institutional and ethical restrictions.

