## Appendix for "Heartbeat-Related Bodily Processing Shapes Transition Patterns in Self-Related Spontaneous Thought"

Original questionnaire

Participants responded to the following questions: (1) During the past week, how often did specific episodes or images spontaneously come to mind? (1. Never, 2. Sometimes (several times per day), 3. Frequently (about once every few hours), 4. Constantly (about once every few minutes)), (2) Which type of episodes or images were most prevalent in your mind during the past week? (1. Personal past experiences, 2. Personal future plans, 3. Knowledge or facts from conversations with others, TV, radio, social media, newspapers, or books, 4. Images or fantasies related to books, videos, music, or artwork that you experienced), (3) Regarding the episodes or images reported in (2), rate the emotional valence and arousal level of the episodes or images themselves (1. Very negative to 5. Very positive / 1. Very calm to 5. Very excited), (4) Regarding the episodes or images reported in (2), rate the emotional valence and arousal level of yourself while recalling these episodes or images (1. Very negative to 5. Very positive / 1. Very calm to 5. Very excited), and (5) To what extent were the episodes or images reported in (2) recalled during the experimental task? (1. Not at all, 2. Sometimes (2-5 times), 3. Frequently (6-9 times), 4. Constantly (10 times or more)). If participants selected "Never" for question (1), the subsequent questions (2-5) were not presented.

Table A1
Model coefficient estimates for differences among Thought Groups in Probe 1 (task concentration) in the asynch condition

| Parameters | Estimate | Est. error | l-95%CI | u-95%CI | Rhat | Bulk_ESS | Tail_ESS |
|---|---|---|---|---|---|---|---|
| Intercept | 63.394 | 3.715 | 56.112 | 70.871 | 1 | 960.342 | 1455.665 |
| Group2 | -19.101 | 2.330 | -23.672 | -14.448 | 1 | 4335.590 | 3030.257 |
| Group3 | -23.185 | 2.274 | -27.682 | -18.820 | 1 | 4552.886 | 3112.075 |
| Group4 | -22.873 | 2.822 | -28.555 | -17.475 | 1 | 4848.202 | 2539.460 |

Table A2
Model coefficient estimates for differences among Thought Groups in Probe 3 (arousal) in the asynch condition

| Parameters | Estimate | Est. error | l-95%CI | u-95%CI | Rhat | Bulk_ESS | Tail_ESS |
|---|---|---|---|---|---|---|---|
| Intercept | 34.608 | 3.155 | 28.413 | 40.820 | 1 | 673.850 | 1404.967 |
| Group2 | -1.081 | 2.696 | -6.313 | 4.331 | 1 | 3560.747 | 2850.601 |
| Group3 | 2.565 | 2.556 | -2.443 | 7.621 | 1 | 3523.281 | 2994.330 |
| Group4 | 1.166 | 3.128 | -4.867 | 7.256 | 1 | 3691.113 | 2885.239 |

Table A3
Model coefficient estimates for differences among Thought Groups in Probe 5 (self-relatedness) in the asynch condition

| Parameters | Estimate | Est. error | l-95%CI | u-95%CI | Rhat | Bulk_ESS | Tail_ESS |
|---|---|---|---|---|---|---|---|
| Intercept | 37.468 | 3.140 | 31.075 | 43.437 | 1 | 737.805 | 1354.876 |
| Group2 | 24.720 | 2.810 | 19.352 | 30.104 | 1 | 3289.510 | 2901.338 |
| Group3 | 29.256 | 2.711 | 24.007 | 34.580 | 1 | 3364.593 | 3330.256 |
| Group4 | -39.009 | 3.255 | -45.388 | -32.587 | 1 | 3151.710 | 3097.732 |

Table A4
Model coefficient estimates for differences among Thought Groups in Probe 7 (bodily information) in the asynch condition

| Parameters | Estimate | Est. error | l-95%CI | u-95%CI | Rhat | Bulk_ESS | Tail_ESS |
|---|---|---|---|---|---|---|---|
| Intercept | 38.487 | 2.944 | 32.827 | 44.280 | 1 | 1716.820 | 2182.330 |
| Group2 | 39.737 | 3.143 | 33.555 | 45.818 | 1 | 3958.124 | 3208.073 |
| Group3 | -10.629 | 3.038 | -16.545 | -4.738 | 1 | 4013.794 | 3200.196 |
| Group4 | -35.663 | 3.692 | -43.031 | -28.259 | 1 | 3722.344 | 3259.639 |

Table A5
Model coefficient estimates for differences among Thought Groups in Probe 8 (contemplation) in the asynch condition

| Parameters | Estimate | Est. error | l-95%CI | u-95%CI | Rhat | Bulk_ESS | Tail_ESS |
|---|---|---|---|---|---|---|---|
| Intercept | 38.121 | 3.315 | 31.481 | 44.429 | 1 | 696.321 | 1522.734 |
| Group2 | 5.678 | 2.813 | 0.235 | 11.122 | 1 | 3624.237 | 3276.330 |
| Group3 | 11.404 | 2.693 | 6.160 | 16.688 | 1 | 3801.543 | 3028.099 |
| Group4 | 3.503 | 3.218 | -2.849 | 9.904 | 1 | 3749.144 | 2735.094 |

Table A6
Model coefficient estimates for differences among Thought Groups in Probe 1 (task concentration) in the synch condition

| Parameters | Estimate | Est. error | l-95%CI | u-95%CI | Rhat | Bulk_ESS | Tail_ESS |
|---|---|---|---|---|---|---|---|
| Intercept | 64.211 | 3.533 | 57.231 | 70.940 | 1 | 1056.253 | 1291.246 |
| Group2 | -18.905 | 2.235 | -23.377 | -14.709 | 1 | 4715.017 | 3039.319 |
| Group3 | -20.207 | 2.324 | -24.825 | -15.638 | 1 | 4430.698 | 2680.211 |
| Group4 | -25.379 | 3.053 | -31.284 | -19.449 | 1 | 4337.239 | 3139.938 |

Table A7
Model coefficient estimates for differences among Thought Groups in Probe 3 (arousal) in the synch condition

| Parameters | Estimate | Est. error | l-95%CI | u-95%CI | Rhat | Bulk_ESS | Tail_ESS |
|---|---|---|---|---|---|---|---|
| Intercept | 29.464 | 2.770 | 24.077 | 35.052 | 1 | 802.527 | 1015.660 |
| Group2 | 4.718 | 2.151 | 0.536 | 8.927 | 1 | 3850.625 | 3091.049 |
| Group3 | 8.118 | 2.215 | 3.897 | 12.383 | 1 | 3792.097 | 2828.609 |
| Group4 | 5.664 | 2.911 | -0.103 | 11.495 | 1 | 3774.500 | 2738.267 |

Table A8
Model coefficient estimates for differences among Thought Groups in Probe 5 (self-relatedness) in the synch condition

| Parameters | Estimate | Est. error | l-95%CI | u-95%CI | Rhat | Bulk_ESS | Tail_ESS |
|---|---|---|---|---|---|---|---|
| Intercept | 36.919 | 2.723 | 31.598 | 42.284 | 1 | 854.198 | 1746.001 |
| Group2 | 19.809 | 2.573 | 14.690 | 24.785 | 1 | 4178.618 | 3290.708 |
| Group3 | 33.093 | 2.687 | 27.978 | 38.429 | 1 | 3696.138 | 2912.415 |
| Group4 | -32.809 | 3.495 | -39.782 | -26.114 | 1 | 3821.718 | 2827.957 |

Table A9
Model coefficient estimates for differences among Thought Groups in Probe 7 (bodily information) in the synch condition

| Parameters | Estimate | Est. error | l-95%CI | u-95%CI | Rhat | Bulk_ESS | Tail_ESS |
|---|---|---|---|---|---|---|---|
| Intercept | 34.840 | 2.680 | 29.582 | 40.014 | 1 | 1798.675 | 2482.182 |
| Group2 | 35.028 | 3.003 | 29.223 | 40.839 | 1 | 3957.557 | 3238.546 |
| Group3 | -9.734 | 3.096 | -15.995 | -3.637 | 1 | 3647.822 | 3420.639 |
| Group4 | -30.681 | 4.022 | -38.824 | -22.740 | 1 | 4270.707 | 3249.970 |

Table A10
Model coefficient estimates for differences among Thought Groups in Probe 8 (contemplation) in the synch condition

| Parameters | Estimate | Est. error | l-95%CI | u-95%CI | Rhat | Bulk_ESS | Tail_ESS |
|---|---|---|---|---|---|---|---|
| Intercept | 35.947 | 3.332 | 29.520 | 42.618 | 1 | 622.091 | 1288.378 |
| Group2 | 6.411 | 2.293 | 1.867 | 10.914 | 1 | 4462.781 | 3221.755 |
| Group3 | 8.200 | 2.392 | 3.512 | 12.773 | 1 | 4656.236 | 3074.491 |
| Group4 | 9.307 | 3.099 | 3.137 | 15.504 | 1 | 4040.305 | 2900.191 |

Table A11
Model coefficient estimates for the Level × Thought Group interaction in Probe 2 (temporal orientation) in the asynch condition

| Parameters | Estimate | Est. error | l-95%CI | u-95%CI | Rhat | Bulk_ESS | Tail_ESS |
|---|---|---|---|---|---|---|---|
| Intercept | 0.001 | 0.028 | -0.053 | 0.057 | 1 | 718.665 | 1448.148 |
| Group2 | -0.001 | 0.043 | -0.085 | 0.086 | 1 | 875.574 | 1618.940 |
| Group3 | 0.293 | 0.042 | 0.213 | 0.378 | 1 | 799.090 | 1598.231 |
| Group4 | 0.126 | 0.047 | 0.037 | 0.219 | 1 | 890.155 | 1597.676 |
| Level2 | 0.000 | 0.040 | -0.079 | 0.078 | 1 | 1010.595 | 1734.651 |
| Level3 | 0.998 | 0.041 | 0.918 | 1.079 | 1 | 934.165 | 1989.786 |
| Level4 | -0.001 | 0.041 | -0.082 | 0.080 | 1 | 1057.891 | 2003.128 |
| Level5 | 0.000 | 0.040 | -0.079 | 0.078 | 1 | 939.978 | 1699.355 |
| Level6 | -0.001 | 0.040 | -0.081 | 0.078 | 1 | 976.554 | 2116.627 |
| Group2:Level2 | 0.000 | 0.060 | -0.120 | 0.123 | 1 | 1238.149 | 2281.605 |
| Group2:Level3 | -0.209 | 0.058 | -0.321 | -0.094 | 1 | 1138.015 | 1859.144 |
| Group2:Level4 | -0.080 | 0.066 | -0.212 | 0.046 | 1 | 1143.224 | 2173.305 |
| Group2:Level5 | 0.002 | 0.060 | -0.118 | 0.121 | 1 | 1187.290 | 2055.098 |
| Group2:Level6 | -1.140 | 0.060 | -1.253 | -1.021 | 1 | 1033.414 | 1808.630 |
| Group3:Level2 | -0.988 | 0.067 | -1.117 | -0.858 | 1 | 1207.313 | 2077.800 |
| Group3:Level3 | 0.000 | 0.061 | -0.119 | 0.119 | 1 | 1252.616 | 1985.958 |
| Group3:Level4 | -0.094 | 0.060 | -0.213 | 0.022 | 1 | 1188.165 | 1989.327 |
| Group3:Level5 | -0.071 | 0.066 | -0.198 | 0.061 | 1 | 1205.753 | 2030.444 |

| | | | | | | | |
|---|---|---|---|---|---|---|---|
| Group3:Level6 | 0.000 | 0.062 | -0.122 | 0.121 | 1 | 1142.616 | 1904.672 |
| Group4:Level2 | -0.176 | 0.059 | -0.291 | -0.059 | 1 | 1111.466 | 2201.412 |
| Group4:Level3 | -0.113 | 0.067 | -0.246 | 0.016 | 1 | 1106.044 | 2100.573 |
| Group4:Level4 | 0.002 | 0.060 | -0.115 | 0.119 | 1 | 1257.626 | 2096.444 |
| Group4:Level5 | -0.139 | 0.059 | -0.253 | -0.022 | 1 | 1081.930 | 1980.784 |
| Group4:Level6 | 0.494 | 0.067 | 0.360 | 0.624 | 1 | 1181.695 | 2045.206 |

Table A12
Model coefficient estimates for the Level × Thought Group interaction in Probe 4 (valence) in the asynch condition

| Parameters | Estimate | Est. error | l-95%CI | u-95%CI | Rhat | Bulk_ESS | Tail_ESS |
|---|---|---|---|---|---|---|---|
| Intercept | 0.010 | 0.040 | -0.066 | 0.089 | 1 | 941.532 | 1700.927 |
| Group2 | 0.097 | 0.060 | -0.019 | 0.216 | 1 | 1051.576 | 1812.577 |
| Group3 | -0.005 | 0.059 | -0.117 | 0.110 | 1 | 1098.004 | 1936.309 |
| Group4 | -0.001 | 0.065 | -0.125 | 0.130 | 1 | 1140.397 | 1982.209 |
| Level2 | 0.179 | 0.056 | 0.068 | 0.289 | 1 | 1188.264 | 1816.230 |
| Level3 | 0.635 | 0.056 | 0.527 | 0.746 | 1 | 1154.417 | 2249.213 |
| Level4 | 0.131 | 0.055 | 0.024 | 0.242 | 1 | 1155.958 | 2240.389 |
| Level5 | -0.002 | 0.057 | -0.111 | 0.110 | 1 | 1237.756 | 2265.428 |
| Group2:Level2 | 0.217 | 0.085 | 0.053 | 0.381 | 1 | 1349.023 | 2186.127 |
| Group2:Level3 | -0.041 | 0.083 | -0.206 | 0.122 | 1 | 1482.948 | 2496.143 |
| Group2:Level4 | -0.008 | 0.090 | -0.190 | 0.167 | 1 | 1427.557 | 2289.074 |
| Group2:Level5 | -0.386 | 0.085 | -0.552 | -0.225 | 1 | 1203.483 | 2062.936 |
| Group3:Level2 | -0.241 | 0.083 | -0.405 | -0.083 | 1 | 1260.485 | 2237.140 |
| Group3:Level3 | -0.129 | 0.092 | -0.307 | 0.052 | 1 | 1454.880 | 2186.655 |
| Group3:Level4 | -0.207 | 0.084 | -0.375 | -0.047 | 1 | 1347.799 | 2527.501 |
| Group3:Level5 | 0.215 | 0.083 | 0.053 | 0.378 | 1 | 1300.694 | 2179.532 |
| Group4:Level2 | 0.083 | 0.092 | -0.097 | 0.259 | 1 | 1602.969 | 2459.405 |
| Group4:Level3 | -0.104 | 0.085 | -0.272 | 0.060 | 1 | 1435.541 | 2238.304 |
| Group4:Level4 | 0.091 | 0.085 | -0.076 | 0.258 | 1 | 1446.108 | 2081.657 |
| Group4:Level5 | 0.052 | 0.092 | -0.131 | 0.226 | 1 | 1499.301 | 2560.388 |

Table A13
Model coefficient estimates for the Level × Thought Group interaction in Probe 6 (perspective) in the asynch condition

| Parameters | Estimate | Est. error | l-95%CI | u-95%CI | Rhat | Bulk_ESS | Tail_ESS |
|---|---|---|---|---|---|---|---|
| Intercept | 0.626 | 0.049 | 0.530 | 0.722 | 1 | 1725.993 | 2467.453 |
| Group2 | 0.275 | 0.075 | 0.126 | 0.419 | 1 | 2437.834 | 2592.224 |
| Group3 | 0.124 | 0.072 | -0.014 | 0.266 | 1 | 2216.367 | 2886.357 |

| | | | | | | | |
|---|---|---|---|---|---|---|---|
| Group4 | -0.146 | 0.081 | -0.304 | 0.012 | 1 | 2071.514 | 2480.046 |
| Level2 | -0.514 | 0.069 | -0.652 | -0.381 | 1 | 1999.684 | 2986.608 |
| Level3 | -0.364 | 0.070 | -0.503 | -0.235 | 1 | 2019.208 | 2891.002 |
| Group2:Level2 | -0.374 | 0.104 | -0.577 | -0.164 | 1 | 2563.972 | 2744.163 |
| Group2:Level3 | -0.093 | 0.101 | -0.290 | 0.108 | 1 | 2527.631 | 2846.580 |
| Group3:Level2 | 0.332 | 0.114 | 0.112 | 0.554 | 1 | 2185.796 | 2971.781 |
| Group3:Level3 | -0.453 | 0.105 | -0.655 | -0.243 | 1 | 2665.497 | 3087.420 |
| Group4:Level2 | -0.287 | 0.102 | -0.482 | -0.086 | 1 | 2620.083 | 3079.702 |
| Group4:Level3 | 0.107 | 0.113 | -0.111 | 0.333 | 1 | 2516.846 | 2730.867 |

Table A14
Model coefficient estimates for the Level × Thought Group interaction in Probe 9 (intentionality) in the asynch condition

| Parameters | Estimate | Est. error | l-95%CI | u-95%CI | Rhat | Bulk_ESS | Tail_ESS |
|---|---|---|---|---|---|---|---|
| Intercept | 0.236 | 0.049 | 0.138 | 0.330 | 1 | 1120.454 | 1750.520 |
| Group2 | 0.074 | 0.073 | -0.066 | 0.218 | 1 | 1326.724 | 1949.990 |
| Group3 | -0.002 | 0.073 | -0.142 | 0.143 | 1 | 1258.438 | 2002.582 |
| Group4 | -0.062 | 0.078 | -0.218 | 0.092 | 1 | 1311.858 | 2211.371 |
| Level2 | 0.172 | 0.068 | 0.039 | 0.308 | 1 | 1238.114 | 1771.026 |
| Level3 | 0.026 | 0.068 | -0.109 | 0.158 | 1 | 1355.824 | 2356.063 |
| Level4 | -0.139 | 0.068 | -0.271 | -0.003 | 1 | 1251.118 | 1884.094 |
| Group2:Level2 | 0.015 | 0.103 | -0.188 | 0.213 | 1 | 1471.101 | 2257.853 |
| Group2:Level3 | 0.148 | 0.102 | -0.057 | 0.350 | 1 | 1365.818 | 2298.581 |
| Group2:Level4 | 0.319 | 0.111 | 0.102 | 0.529 | 1 | 1645.740 | 2398.430 |
| Group3:Level2 | -0.181 | 0.101 | -0.382 | 0.015 | 1 | 1650.700 | 2264.797 |
| Group3:Level3 | -0.086 | 0.101 | -0.289 | 0.111 | 1 | 1532.939 | 2208.946 |
| Group3:Level4 | -0.064 | 0.111 | -0.273 | 0.157 | 1 | 1609.082 | 2492.773 |
| Group4:Level2 | -0.128 | 0.100 | -0.328 | 0.070 | 1 | 1641.083 | 2363.093 |
| Group4:Level3 | -0.058 | 0.102 | -0.254 | 0.143 | 1 | 1491.735 | 2478.242 |
| Group4:Level4 | -0.005 | 0.110 | -0.217 | 0.209 | 1 | 1518.769 | 2573.835 |

Table A15
Model coefficient estimates for the Level × Thought Group interaction in Probe 2 (temporal orientation) in the synch condition

| Parameters | Estimate | Est. error | l-95%CI | u-95%CI | Rhat | Bulk_ESS | Tail_ESS |
|---|---|---|---|---|---|---|---|
| Intercept | 0.000 | 0.028 | -0.054 | 0.056 | 1 | 610.660 | 1423.017 |
| Group2 | 0.000 | 0.040 | -0.079 | 0.078 | 1 | 804.896 | 1651.202 |
| Group3 | 0.192 | 0.041 | 0.108 | 0.272 | 1 | 713.757 | 1624.010 |

| | | | | | | | |
|---|---|---|---|---|---|---|---|
| Group4 | 0.055 | 0.046 | -0.035 | 0.143 | 1 | 835.770 | 1736.713 |
| Level2 | 0.000 | 0.040 | -0.078 | 0.078 | 1 | 884.370 | 1808.502 |
| Level3 | 1.000 | 0.039 | 0.923 | 1.076 | 1 | 881.066 | 1802.949 |
| Level4 | 0.001 | 0.040 | -0.078 | 0.076 | 1 | 862.706 | 2052.809 |
| Level5 | 0.000 | 0.039 | -0.078 | 0.077 | 1 | 943.610 | 1947.170 |
| Level6 | 0.000 | 0.040 | -0.076 | 0.079 | 1 | 845.557 | 1909.581 |
| Group2:Level2 | -0.001 | 0.057 | -0.111 | 0.116 | 1 | 1179.278 | 2138.672 |
| Group2:Level3 | -0.011 | 0.058 | -0.123 | 0.105 | 1 | 1003.074 | 2215.453 |
| Group2:Level4 | 0.028 | 0.065 | -0.099 | 0.155 | 1 | 1196.910 | 2634.619 |
| Group2:Level5 | 0.000 | 0.057 | -0.111 | 0.112 | 1 | 1097.257 | 2014.543 |
| Group2:Level6 | -1.051 | 0.057 | -1.160 | -0.939 | 1 | 1032.347 | 2219.875 |
| Group3:Level2 | -0.864 | 0.065 | -0.990 | -0.737 | 1 | 1280.751 | 2564.598 |
| Group3:Level3 | -0.001 | 0.058 | -0.115 | 0.114 | 1 | 1152.903 | 2407.251 |
| Group3:Level4 | 0.020 | 0.058 | -0.092 | 0.134 | 1 | 995.752 | 2056.816 |
| Group3:Level5 | -0.034 | 0.065 | -0.159 | 0.094 | 1 | 1178.299 | 2461.933 |
| Group3:Level6 | 0.000 | 0.057 | -0.112 | 0.112 | 1 | 1187.445 | 2200.955 |
| Group4:Level2 | -0.086 | 0.058 | -0.198 | 0.030 | 1 | 1103.775 | 1760.254 |
| Group4:Level3 | 0.006 | 0.064 | -0.117 | 0.135 | 1 | 1179.269 | 2466.292 |
| Group4:Level4 | -0.001 | 0.057 | -0.111 | 0.110 | 1 | 1112.481 | 2092.087 |
| Group4:Level5 | -0.043 | 0.057 | -0.155 | 0.069 | 1 | 964.050 | 1929.662 |
| Group4:Level6 | 0.491 | 0.064 | 0.363 | 0.615 | 1 | 1126.145 | 2301.916 |

Table A16
Model coefficient estimates for the Level × Thought Group interaction in Probe 4 (valence) in the synch condition

| Parameters | Estimate | Est. error | l-95%CI | u-95%CI | Rhat | Bulk_ESS | Tail_ESS |
|---|---|---|---|---|---|---|---|
| Intercept | 0.017 | 0.038 | -0.056 | 0.090 | 1 | 820.374 | 1788.498 |
| Group2 | 0.054 | 0.056 | -0.056 | 0.165 | 1 | 1042.234 | 1659.357 |
| Group3 | 0.003 | 0.056 | -0.107 | 0.109 | 1 | 950.601 | 1952.834 |
| Group4 | -0.005 | 0.064 | -0.127 | 0.119 | 1 | 1090.464 | 2073.253 |
| Level2 | 0.099 | 0.055 | -0.012 | 0.206 | 1 | 1122.080 | 1937.438 |
| Level3 | 0.754 | 0.055 | 0.651 | 0.863 | 1 | 1083.028 | 2381.795 |
| Level4 | 0.073 | 0.054 | -0.033 | 0.179 | 1 | 1019.108 | 1988.657 |
| Level5 | -0.013 | 0.055 | -0.123 | 0.093 | 1 | 1140.069 | 2277.374 |
| Group2:Level2 | 0.278 | 0.080 | 0.122 | 0.439 | 1 | 1458.186 | 2369.491 |
| Group2:Level3 | 0.109 | 0.080 | -0.051 | 0.263 | 1 | 1269.787 | 2497.002 |
| Group2:Level4 | 0.132 | 0.089 | -0.040 | 0.303 | 1 | 1461.837 | 2484.719 |

| | | | | | | | |
|---|---|---|---|---|---|---|---|
| Group2:Level5 | -0.407 | 0.079 | -0.565 | -0.250 | 1 | 1410.329 | 2516.337 |
| Group3:Level2 | -0.408 | 0.079 | -0.562 | -0.258 | 1 | 1242.186 | 2293.147 |
| Group3:Level3 | -0.201 | 0.090 | -0.380 | -0.027 | 1 | 1402.335 | 2553.605 |
| Group3:Level4 | -0.090 | 0.077 | -0.238 | 0.071 | 1 | 1304.230 | 1985.482 |
| Group3:Level5 | 0.266 | 0.080 | 0.112 | 0.421 | 1 | 1252.707 | 2257.656 |
| Group4:Level2 | 0.075 | 0.089 | -0.106 | 0.241 | 1 | 1384.628 | 2561.861 |
| Group4:Level3 | -0.049 | 0.080 | -0.209 | 0.109 | 1 | 1338.146 | 2457.746 |
| Group4:Level4 | 0.024 | 0.080 | -0.133 | 0.179 | 1 | 1302.651 | 2484.561 |
| Group4:Level5 | 0.022 | 0.090 | -0.156 | 0.202 | 1 | 1352.093 | 2362.304 |

Table A17
Model coefficient estimates for the Level × Thought Group interaction in Probe 6 (perspective) in the synch condition

| Parameters | Estimate | Est. error | l-95%CI | u-95%CI | Rhat | Bulk_ESS | Tail_ESS |
|---|---|---|---|---|---|---|---|
| Intercept | 0.688 | 0.046 | 0.599 | 0.778 | 1 | 1457.304 | 2510.244 |
| Group2 | 0.228 | 0.066 | 0.098 | 0.353 | 1 | 1844.347 | 2512.818 |
| Group3 | 0.090 | 0.066 | -0.038 | 0.218 | 1 | 1792.111 | 2538.675 |
| Group4 | -0.211 | 0.074 | -0.355 | -0.070 | 1 | 2009.117 | 2028.548 |
| Level2 | -0.633 | 0.065 | -0.762 | -0.505 | 1 | 1547.060 | 2463.572 |
| Level3 | -0.442 | 0.064 | -0.567 | -0.316 | 1 | 1632.886 | 2413.787 |
| Group2:Level2 | -0.273 | 0.093 | -0.456 | -0.089 | 1 | 2005.764 | 2354.430 |
| Group2:Level3 | 0.003 | 0.094 | -0.181 | 0.190 | 1 | 2096.973 | 2706.439 |
| Group3:Level2 | 0.411 | 0.106 | 0.207 | 0.628 | 1 | 2163.440 | 2645.337 |
| Group3:Level3 | -0.407 | 0.093 | -0.586 | -0.223 | 1 | 2030.522 | 2931.298 |
| Group4:Level2 | -0.262 | 0.093 | -0.444 | -0.078 | 1 | 2091.962 | 3167.535 |
| Group4:Level3 | 0.232 | 0.103 | 0.031 | 0.433 | 1 | 2141.483 | 2769.638 |

Table A18
Model coefficient estimates for the Level × Thought Group interaction in Probe 9 (intentionality) in the synch condition

| Parameters | Estimate | Est. error | l-95%CI | u-95%CI | Rhat | Bulk_ESS | Tail_ESS |
|---|---|---|---|---|---|---|---|
| Intercept | 0.155 | 0.046 | 0.068 | 0.246 | 1 | 952.012 | 1498.864 |
| Group2 | 0.070 | 0.067 | -0.067 | 0.201 | 1 | 1273.145 | 1938.588 |
| Group3 | 0.065 | 0.067 | -0.070 | 0.194 | 1 | 1189.219 | 1755.022 |
| Group4 | 0.070 | 0.077 | -0.078 | 0.219 | 1 | 1189.965 | 2496.266 |
| Level2 | 0.307 | 0.067 | 0.176 | 0.436 | 1 | 1218.643 | 1908.687 |
| Level3 | 0.147 | 0.066 | 0.017 | 0.276 | 1 | 1260.723 | 2228.881 |
| Level4 | -0.076 | 0.065 | -0.208 | 0.047 | 1 | 1355.279 | 1921.334 |

| | | | | | | | |
|---|---|---|---|---|---|---|---|
| Group2:Level2 | 0.048 | 0.097 | -0.139 | 0.242 | 1 | 1466.093 | 1829.589 |
| Group2:Level3 | 0.037 | 0.099 | -0.155 | 0.230 | 1 | 1403.504 | 2182.049 |
| Group2:Level4 | 0.068 | 0.108 | -0.150 | 0.280 | 1 | 1593.386 | 2539.212 |
| Group3:Level2 | -0.217 | 0.096 | -0.398 | -0.025 | 1 | 1675.057 | 2306.693 |
| Group3:Level3 | -0.169 | 0.096 | -0.355 | 0.023 | 1 | 1454.240 | 2256.599 |
| Group3:Level4 | -0.219 | 0.111 | -0.439 | -0.002 | 1 | 1673.313 | 2908.111 |
| Group4:Level2 | -0.122 | 0.096 | -0.305 | 0.071 | 1 | 1614.659 | 2361.993 |
| Group4:Level3 | -0.124 | 0.095 | -0.307 | 0.061 | 1 | 1615.361 | 2393.923 |
| Group4:Level4 | -0.132 | 0.109 | -0.344 | 0.087 | 1 | 1747.296 | 2630.979 |

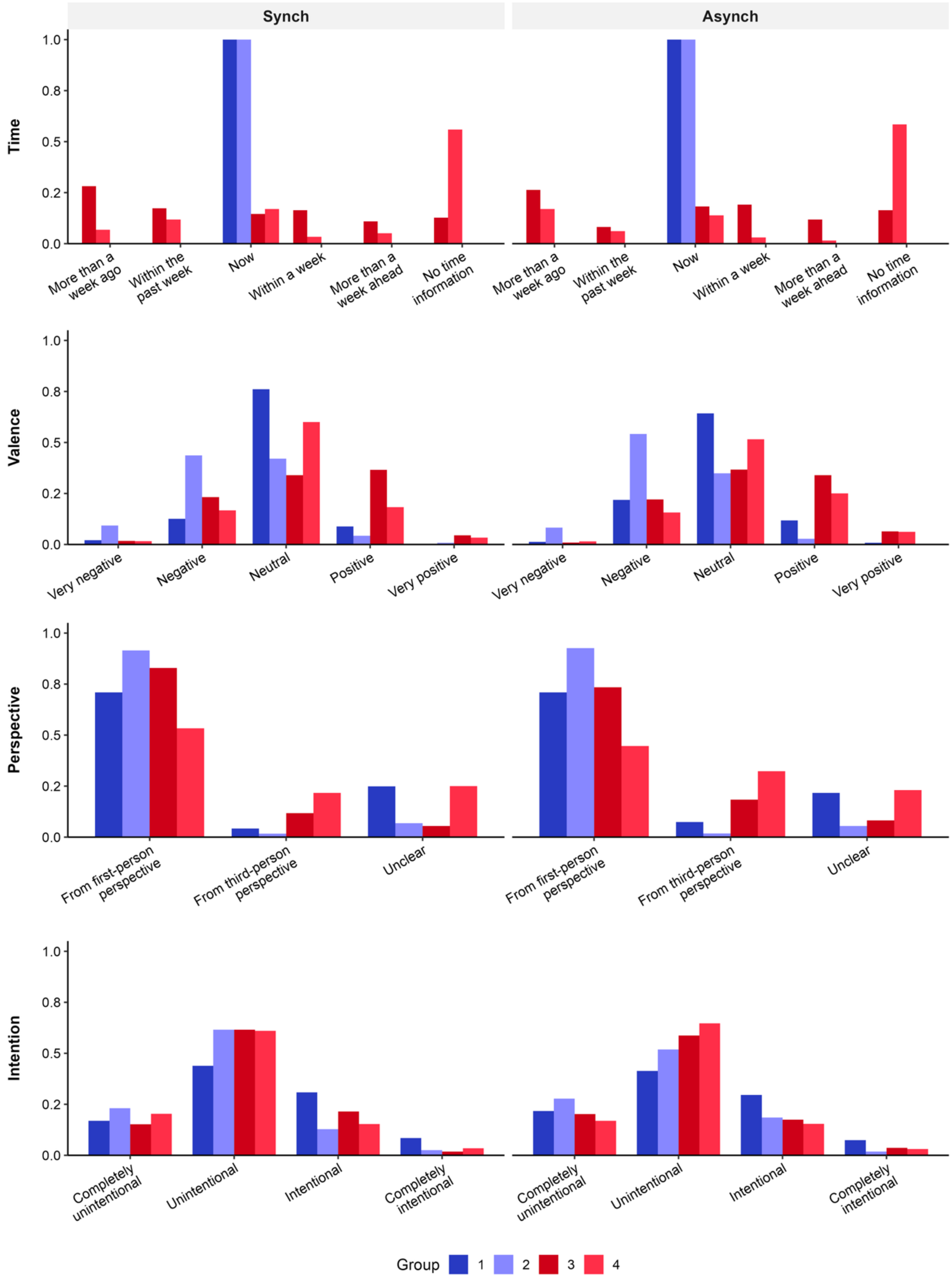

Synch
Asynch
Time
Valence
Perspective
Intention
1.0
0.8
0.5
0.2
0.0
More than a week ago
Within the past week
Now
Within a week
More than a week ahead
No time information
Very negative
Negative
Neutral
Positive
Very positive
From first-person perspective
From third-person perspective
Unclear
Completely unintentional
Unintentional
Intentional
Completely intentional
Group
1
2
3
4


Figure A1

Report proportions of discrete probe levels for each thought group across experimental conditions. We examined how the categorized thought groups—defined by task-relatedness and stimulus-independence—reflected the qualitative dimensions of reported thoughts (Time, Valence, Perspective, and Intention). In both the synch and asynch conditions, the Task-focus state (Group 1) was characterized by a dominance of "now" and "neutral" thoughts. In contrast, internal thought states exhibited distinct qualitative profiles: Group 2 (Stimulus-dependent) showed higher tendencies toward "negative" and "intentional" reports, while Group 3 (Self-related) was associated with an increase in "from third-person perspective" and "very positive" ratings. These results demonstrate the unique qualitative properties inherent to each thought group.

Table A19
Bayesian mixed-effects model of reaction-time variability across experimental conditions and thought groups.

| Parameters | Estimate | SE | 95% CI | Rhat | Bulk_ESS | Tail_ESS |
|---|---|---|---|---|---|---|
| Intercept | 0.283 | 0.029 | [0.226, 0.340] | 1 | 2476 | 4901 |
| Condition | -0.045 | 0.028 | [-0.101, 0.010] | 1 | 4374 | 7765 |
| Group 2 | 0.01 | 0.025 | [-0.038, 0.058] | 1 | 8068 | 9126 |
| Group 3 | -0.025 | 0.024 | [-0.072, 0.021] | 1 | 7970 | 8517 |
| Group 4 | -0.009 | 0.027 | [-0.062, 0.044] | 1 | 8480 | 8723 |
| Group 5 | 0.004 | 0.026 | [-0.047, 0.054] | 1 | 8487 | 9808 |
| Condition: Group 2 | 0.02 | 0.034 | [-0.045, 0.088] | 1 | 7319 | 8662 |
| Condition: Group 3 | 0.026 | 0.033 | [-0.040, 0.092] | 1 | 7427 | 8221 |
| Condition: Group 4 | 0.047 | 0.038 | [-0.028, 0.124] | 1 | 7878 | 9115 |
| Condition: Group 5 | 0.036 | 0.037 | [-0.037, 0.107] | 1 | 7572 | 9100 |

Table A20
Bayesian binomial mixed-effects model of response error rate across experimental conditions and thought groups.

| Parameters | Estimate | SE | 95% CI | Rhat | Bulk_ESS | Tail_ESS |
|---|---|---|---|---|---|---|
| Intercept | -3.826 | 0.251 | [-4.324, -3.343] | 1 | 1252 | 2188 |
| Condition | -1.199 | 0.303 | [-1.818, -0.633] | 1 | 2433 | 4735 |
| Group 2 | -0.66 | 0.098 | [-0.854, -0.470] | 1 | 9209 | 9168 |
| Group 3 | -0.611 | 0.092 | [-0.792, -0.430] | 1 | 9292 | 9694 |
| Group 4 | -0.566 | 0.091 | [-0.746, -0.390] | 1 | 10031 | 9192 |
| Group 5 | -0.388 | 0.095 | [-0.577, -0.201] | 1 | 9097 | 8822 |
| Condition: Group 2 | 0.461 | 0.151 | [0.167, 0.756] | 1 | 9125 | 8967 |
| Condition: Group 3 | 0.016 | 0.146 | [-0.274, 0.301] | 1 | 9381 | 9262 |
| Condition: Group 4 | 0.394 | 0.133 | [0.133, 0.652] | 1 | 8950 | 9493 |
| Condition: Group 5 | 0.614 | 0.137 | [0.345, 0.884] | 1 | 8454 | 8981 |

Table A21
Posterior synch–asynch contrasts in response error probability within each thought group

| Group | Asynch | Synch | Synch - Asynch | 95% CI | P_direction |
|---|---|---|---|---|---|
| Group 1 | 2.20 | 0.70 | -1.50 | [-2.46, -0.76] | 1.000 |
| Group 2 | 1.15 | 0.58 | -0.57 | [-1.12, -0.12] | 0.992 |
| Group 3 | 1.21 | 0.39 | -0.82 | [-1.39, -0.39] | 1.000 |

| | | | | | |
|---|---|---|---|---|---|
| Group 4 | 1.26 | 0.59 | -0.67 | [-1.27, -0.19] | 0.995 |
| Group 5 | 1.51 | 0.88 | -0.63 | [-1.35, 0.02] | 0.972 |

Table A22
Bayesian mixed-effects model of standardized RR intervals across experimental conditions and thought groups.

| Parameters | Estimate | SE | 95% CI | Rhat | Bulk_ESS | Tail_ESS |
|---|---|---|---|---|---|---|
| Intercept | 0.108 | 0.036 | [0.038, 0.180] | 1 | 8645 | 8933 |
| Condition | -0.057 | 0.05 | [-0.157, 0.043] | 1 | 8000 | 7995 |
| Group 2 | -0.121 | 0.063 | [-0.246, -0.001] | 1 | 9486 | 8339 |
| Group 3 | -0.183 | 0.064 | [-0.309, -0.057] | 1 | 9497 | 9109 |
| Group 4 | -0.15 | 0.077 | [-0.301, -0.000] | 1 | 11538 | 10476 |
| Group 5 | -0.147 | 0.071 | [-0.288, -0.005] | 1 | 9949 | 8195 |
| Condition: Group 2 | 0.123 | 0.088 | [-0.047, 0.295] | 1 | 9374 | 9438 |
| Condition: Group 3 | 0.004 | 0.089 | [-0.169, 0.176] | 1 | 8790 | 9255 |
| Condition: Group 4 | 0.047 | 0.109 | [-0.167, 0.265] | 1 | 10969 | 9042 |
| Condition: Group 5 | 0.037 | 0.104 | [-0.164, 0.240] | 1 | 10561 | 9198 |

Table A23
Posterior contrasts for standardized RR intervals across experimental conditions and thought groups.

| Condition | Contrast | Estimate | 95% CI | P_direction |
|---|---|---|---|---|
| Asynch | Group 2 - Group 1 | -0.121 | [-0.246, -0.001] | 0.976 |
| Asynch | Group 3 - Group 1 | -0.183 | [-0.309, -0.057] | 0.998 |
| Asynch | Group 4 - Group 1 | -0.15 | [-0.301, -0.000] | 0.975 |
| Asynch | Group 5 - Group 1 | -0.147 | [-0.288, -0.005] | 0.979 |
| Synch | Group 2 - Group 1 | 0.002 | [-0.117, 0.124] | 0.514 |
| Synch | Group 3 - Group 1 | -0.179 | [-0.304, -0.055] | 0.998 |
| Synch | Group 4 - Group 1 | -0.103 | [-0.256, 0.053] | 0.904 |
| Synch | Group 5 - Group 1 | -0.11 | [-0.259, 0.043] | 0.925 |

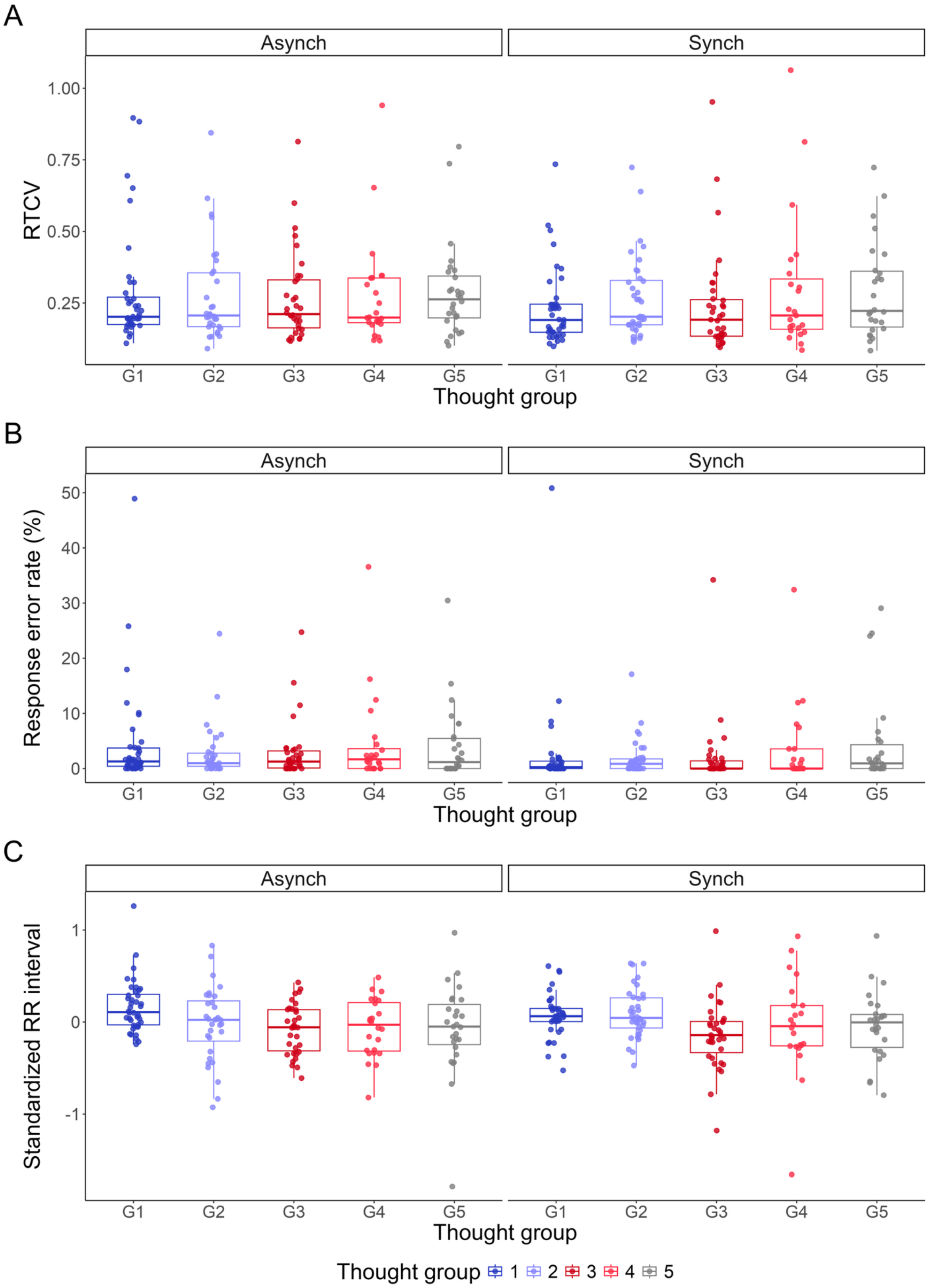
A
Asynch
Synch
RTCV
1.00
0.75
0.50
0.25
G1
G2
G3
G4
G5
Thought group
B
Response error rate (%)
50
40
30
20
10
0
C
Standardized RR interval
1
0
-1
Thought group 1 2 3 4 5

Figure A2

Objective behavioral and cardiac characteristics of thought groups across experimental conditions

The three panels show the distributions of objective measures across the five thought groups separately for the asynch and synch conditions: (A) reaction-time coefficient of variation (RTCV), (B) response error rate, and (C) standardized RR interval. Individual points represent participant-level values for each thought group and condition. Box plots indicate the median and interquartile range, with whiskers extending to 1.5 times the interquartile range. RR intervals were standardized within participant across the task. Colors indicate thought group: Group 1 (On-task), Group 2 (Stimulus-dependent), Group 3 (Self-related), Group 4 (Task/Self-unrelated), and Group 5 (Thought absence).

Table A24

Bayesian multinomial mixed-effects model of thought-group report distribution as a function of condition

| Parameter | Estimate | Est.Error | l-95% CI | u-95% CI | Rhat | Bulk_ESS | Tail_ESS |
|---|---|---|---|---|---|---|---|
| G2_Intercept | -0.883 | 0.178 | -1.246 | -0.546 | 1 | 5282.425 | 7663.905 |
| G3_Intercept | -0.884 | 0.181 | -1.245 | -0.538 | 1 | 5901.654 | 8009.529 |
| G4_Intercept | -1.701 | 0.276 | -2.284 | -1.193 | 1 | 4411.108 | 6927.778 |
| G5_Intercept | -1.416 | 0.258 | -1.953 | -0.943 | 1 | 3922.838 | 6012.079 |
| G2_Synch-Asynch | 0.060 | 0.166 | -0.263 | 0.385 | 1 | 12431.867 | 9397.586 |
| G3_Synch-Asynch | -0.013 | 0.167 | -0.342 | 0.316 | 1 | 14088.143 | 9967.539 |
| G4_Synch-Asynch | -0.125 | 0.210 | -0.538 | 0.287 | 1 | 13058.631 | 8861.171 |
| G5_Synch-Asynch | -0.281 | 0.197 | -0.672 | 0.103 | 1 | 13216.018 | 8522.436 |

Note. Coefficients are on the log-odds scale. On-task thought (G1) and the asynch condition served as the reference outcome category and reference condition, respectively. Intercept terms represent the relative log-odds of each thought group compared with G1 in the asynch condition, whereas Synch–Asynch terms represent condition-related changes in these relative log-odds. G2 = Stimulus-dependent thought; G3 = Self-related thought; G4 = Task/Self-unrelated thought; G5 = Thought absence.

Table A25

Synch–asynch contrasts in transition probabilities between thought groups

| Transition | Estimate | CI_lower | CI_upper | P_gt_0 | P_lt_0 |
|---|---|---|---|---|---|
| G1 → G1 | 0.007 | -0.091 | 0.104 | 0.561 | 0.439 |
| G1 → G2 | 0.053 | -0.017 | 0.127 | 0.932 | 0.068 |
| G1 → G3 | -0.060 | -0.139 | 0.015 | 0.060 | 0.940 |
| G1 → G4 | -0.010 | -0.064 | 0.040 | 0.346 | 0.654 |
| G1 → G5 | 0.010 | -0.038 | 0.060 | 0.655 | 0.345 |
| G2 → G1 | 0.098 | -0.046 | 0.237 | 0.910 | 0.090 |
| G2 → G2 | -0.070 | -0.193 | 0.047 | 0.125 | 0.875 |
| G2 → G3 | 0.081 | -0.042 | 0.205 | 0.906 | 0.094 |
| G2 → G4 | -0.037 | -0.112 | 0.023 | 0.110 | 0.890 |
| G2 → G5 | -0.071 | -0.188 | 0.029 | 0.085 | 0.915 |
| G3 → G1 | -0.022 | -0.167 | 0.126 | 0.382 | 0.618 |
| G3 → G2 | 0.065 | -0.055 | 0.187 | 0.857 | 0.143 |
| G3 → G3 | 0.023 | -0.073 | 0.125 | 0.667 | 0.333 |
| G3 → G4 | 0.013 | -0.057 | 0.089 | 0.638 | 0.362 |
| G3 → G5 | -0.079 | -0.179 | 0.005 | 0.034 | 0.966 |

| | | | | | |
|---|---|---|---|---|---|
| G4 → G1 | 0.043 | -0.144 | 0.233 | 0.668 | 0.332 |
| G4 → G2 | -0.061 | -0.214 | 0.094 | 0.205 | 0.795 |
| G4 → G3 | 0.057 | -0.121 | 0.236 | 0.739 | 0.261 |
| G4 → G4 | 0.019 | -0.076 | 0.119 | 0.659 | 0.341 |
| G4 → G5 | -0.058 | -0.216 | 0.088 | 0.215 | 0.785 |
| G5 → G1 | -0.010 | -0.184 | 0.167 | 0.456 | 0.544 |
| G5 → G2 | 0.027 | -0.147 | 0.201 | 0.617 | 0.383 |
| G5 → G3 | -0.029 | -0.178 | 0.120 | 0.346 | 0.654 |
| G5 → G4 | 0.008 | -0.107 | 0.135 | 0.548 | 0.452 |
| G5 → G5 | 0.003 | -0.069 | 0.079 | 0.534 | 0.466 |

Note. Estimates indicate synch–asynch differences in posterior predicted transition probabilities. Positive values indicate higher probabilities in the synch condition. CI = credible interval. G1 = On-task thought; G2 = Stimulus-dependent thought; G3 = Self-related thought; G4 = Task/Self-unrelated thought; G5 = Thought absence. P_gt_0 indicates the posterior probability that the synch–asynch difference was greater than zero; P_lt_0 indicates the posterior probability that the difference was less than zero.

Table A26
Fixed-effect estimates from the Bayesian multinomial mixed-effects model of thought report distribution

| Term | Outcome | Estimate | Est.Error | l-95% CI | u-95% CI | Rhat | Bulk_ESS | Tail_ESS |
|---|---|---|---|---|---|---|---|---|
| Intercept | G2 vs G1 | -0.876 | 0.187 | -1.252 | -0.515 | 1 | 6072 | 8207 |
| Intercept | G3 vs G1 | -0.887 | 0.182 | -1.254 | -0.536 | 1 | 7200 | 8829 |
| Intercept | G4 vs G1 | -1.792 | 0.298 | -2.420 | -1.243 | 1 | 5606 | 7755 |
| Intercept | G5 vs G1 | -1.507 | 0.282 | -2.101 | -0.985 | 1 | 4476 | 5888 |
| Condition: Synch | G2 vs G1 | 0.027 | 0.173 | -0.311 | 0.367 | 1 | 18259 | 8762 |
| Condition: Synch | G3 vs G1 | 0.029 | 0.176 | -0.320 | 0.373 | 1 | 16984 | 9779 |
| Condition: Synch | G4 vs G1 | -0.195 | 0.231 | -0.657 | 0.254 | 1 | 17701 | 9283 |
| Condition: Synch | G5 vs G1 | -0.270 | 0.208 | -0.680 | 0.141 | 1 | 18054 | 8737 |
| HCT score (z) | G2 vs G1 | 0.126 | 0.184 | -0.232 | 0.485 | 1 | 5759 | 7934 |
| HCT score (z) | G3 vs G1 | -0.200 | 0.186 | -0.560 | 0.169 | 1 | 6475 | 7849 |
| HCT score (z) | G4 vs G1 | 0.206 | 0.290 | -0.372 | 0.767 | 1 | 5257 | 7345 |
| HCT score (z) | G5 vs G1 | 0.253 | 0.272 | -0.273 | 0.803 | 1 | 4532 | 6848 |
| Condition: Synch × HCT score (z) | G2 vs G1 | -0.068 | 0.172 | -0.404 | 0.269 | 1 | 12170 | 9567 |
| Condition: Synch × HCT score (z) | G3 vs G1 | 0.244 | 0.179 | -0.106 | 0.595 | 1 | 12897 | 9456 |
| Condition: Synch × HCT score (z) | G4 vs G1 | -0.793 | 0.234 | -1.261 | -0.336 | 1 | 15841 | 9798 |
| Condition: Synch × HCT score (z) | G5 vs G1 | -0.087 | 0.210 | -0.497 | 0.327 | 1 | 13808 | 9295 |

Table A27
Posterior estimates from the Bayesian linear mixed-effects model of HEP amplitude

| Parameter | Estimate | Est.Error | l-95% CI | u-95% CI | Rhat | Bulk_ESS | Tail_ESS |
|---|---|---|---|---|---|---|---|
| Intercept | -0.962 | 0.108 | -1.174 | -0.749 | 1 | 2627.360 | 4828.963 |
| Group2 | 0.095 | 0.075 | -0.054 | 0.242 | 1 | 12401.092 | 9898.048 |
| Group3 | 0.012 | 0.069 | -0.124 | 0.148 | 1 | 12524.596 | 9881.910 |
| Group4 | 0.095 | 0.086 | -0.074 | 0.262 | 1 | 12004.577 | 9831.170 |

| | | | | | | | |
|---|---|---|---|---|---|---|---|
| Group5 | 0.218 | 0.088 | 0.045 | 0.392 | 1 | 12708.980 | 10343.111 |
| Condition | -0.214 | 0.125 | -0.460 | 0.034 | 1 | 4365.337 | 6500.848 |
| n_trial_z | 0.010 | 0.021 | -0.031 | 0.053 | 1 | 16834.568 | 10551.834 |
| Group2:Condition | -0.084 | 0.103 | -0.288 | 0.120 | 1 | 13445.612 | 9891.814 |
| Group3:Condition | -0.058 | 0.096 | -0.247 | 0.130 | 1 | 13549.350 | 9706.812 |
| Group4:Condition | -0.016 | 0.121 | -0.254 | 0.222 | 1 | 14805.154 | 9991.614 |
| Group5:Condition | -0.171 | 0.127 | -0.419 | 0.081 | 1 | 15412.762 | 9677.301 |

Table A28

Sensitivity of HEP condition effects to the AEP-correction procedure

| AEP correction | ThoughtGroup | Synch - Asynch Estimate (uV) | Synch - Asynch 95% CrI | Difference vs Group 1 Estimate (uV) | Difference vs Group 1 95% CrI |
|---|---|---|---|---|---|
| Uncorrected | Group 1 | -0.046 | [-0.295, 0.206] | | |
| Uncorrected | Group 2 | -0.111 | [-0.392, 0.174] | -0.065 | [-0.279, 0.149] |
| Uncorrected | Group 3 | -0.119 | [-0.390, 0.155] | -0.073 | [-0.273, 0.121] |
| Uncorrected | Group 4 | -0.094 | [-0.401, 0.217] | -0.048 | [-0.294, 0.198] |
| Uncorrected | Group 5 | -0.245 | [-0.565, 0.075] | -0.199 | [-0.461, 0.064] |
| Pooled AEP correction | Group 1 | -0.208 | [-0.456, 0.039] | | |
| Pooled AEP correction | Group 2 | -0.258 | [-0.543, 0.025] | -0.05 | [-0.264, 0.163] |
| Pooled AEP correction | Group 3 | -0.281 | [-0.550, -0.006] | -0.073 | [-0.271, 0.120] |
| Pooled AEP correction | Group 4 | -0.255 | [-0.575, 0.062] | -0.046 | [-0.299, 0.209] |
| Pooled AEP correction | Group 5 | -0.394 | [-0.715, -0.078] | -0.186 | [-0.455, 0.080] |
| Thought-group-specific AEP correction | Group 1 | -0.235 | [-0.502, 0.036] | | |
| Thought-group-specific AEP correction | Group 2 | -0.262 | [-0.582, 0.051] | -0.027 | [-0.293, 0.242] |
| Thought-group-specific AEP correction | Group 3 | -0.285 | [-0.594, 0.016] | -0.05 | [-0.301, 0.196] |
| Thought-group-specific AEP correction | Group 4 | -0.201 | [-0.568, 0.160] | 0.034 | [-0.286, 0.345] |
| Thought-group-specific AEP correction | Group 5 | -0.458 | [-0.807, -0.107] | -0.223 | [-0.530, 0.086] |

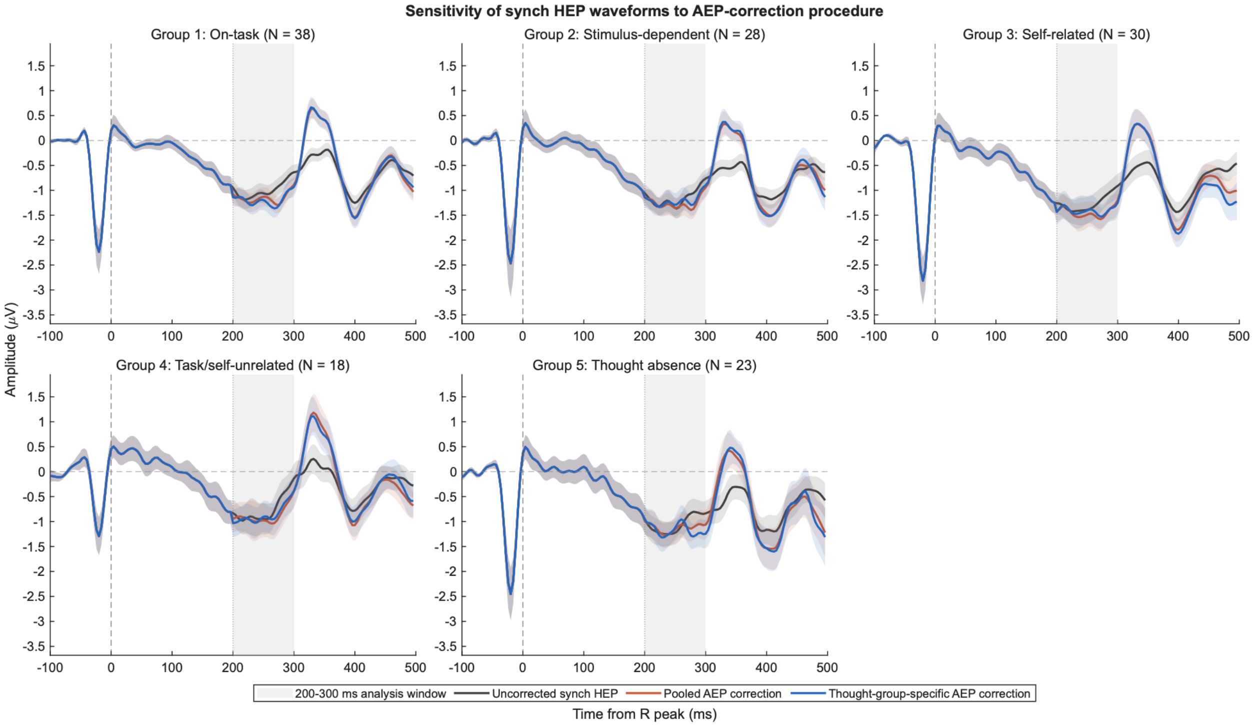


Figure A3
Sensitivity of synch HEP waveforms to the AEP-correction procedure.
Grand-average synch HEP waveforms are shown separately for the five thought groups using the same matched participant sample within each group. Black lines show uncorrected synch HEPs, red lines show HEPs after participant-specific pooled AEP correction, and blue lines show HEPs after participant- and thought-group-specific AEP correction. The gray vertical region indicates the predefined 200–300 ms HEP analysis window, and the vertical dashed line indicates the R peak (0 ms). The number of participants contributing matched estimates is shown above each panel. The corresponding model-based Synch–Asynch contrasts for the three correction procedures are reported in Table A28.

Table 29
Sensitivity of HEP condition effects to the modelling of estimate-specific sampling uncertainty

| Precision model | Thought Group | Synch - Asynch Estimate (uV) | Synch - Asynch 95% CrI | Difference vs Group 1 Estimate (uV) | Difference vs Group 1 95% CrI |
|---|---|---|---|---|---|
| Cell-SE model | Group 1 | -0.214 | [-0.460, 0.034] | | |
| Cell-SE model | Group 2 | -0.298 | [-0.573, -0.019] | -0.084 | [-0.288, 0.120] |
| Cell-SE model | Group 3 | -0.272 | [-0.534, -0.007] | -0.058 | [-0.247, 0.130] |
| Cell-SE model | Group 4 | -0.23 | [-0.526, 0.073] | -0.016 | [-0.254, 0.222] |
| Cell-SE model | Group 5 | -0.385 | [-0.694, -0.074] | -0.171 | [-0.419, 0.081] |
| Model without cell-specific SE | Group 1 | -0.179 | [-0.515, 0.164] | | |
| Model without cell-specific SE | Group 2 | -0.314 | [-0.660, 0.037] | -0.135 | [-0.499, 0.229] |
| Model without cell-specific SE | Group 3 | -0.286 | [-0.633, 0.071] | -0.107 | [-0.467, 0.258] |

| | | | | | |
|---|---|---|---|---|---|
| Model without cell-specific SE | Group 4 | -0.018 | [-0.421, 0.382] | 0.161 | [-0.244, 0.578] |
| Model without cell-specific SE | Group 5 | -0.347 | [-0.724, 0.033] | -0.169 | [-0.563, 0.227] |
